# Motor learning shapes temporal activity in human sensorimotor cortex

**DOI:** 10.1101/345421

**Authors:** Catharina Zich, Mark W Woolrich, Robert Becker, Diego Vidaurre, Jacqueline Scholl, Emily L Hinson, Laurie Josephs, Sven Braeutigam, Andrew J Quinn, Charlotte J Stagg

## Abstract

Although neuroimaging techniques have provided vital insights into the anatomical regions involved in motor learning, the underlying changes in temporal dynamics are not well understood. Using magnetoencephalography and Hidden Markov Modelling to model the dynamics of neural oscillations on data-adaptive time-scales, we detected specific changes in movement-related sensorimotor β-activity during practice of a self-paced sequential visuo-motor task. The behaviourally-relevant neural signature generalised to another motor task, emphasising the centrality of β-activity in motor plasticity.

## Main

Plasticity refers to the ability of organisms or cells to alter their phenotype in response to changes. One fundamental example of human plasticity which may serve as an exemplar is learning a new motor skill. (f)MRI and PET studies have provided critical insights as to which brain regions are involved in motor learning^1,2^. However, the temporal dynamics underpinning motor skill acquisition are not well understood. To date, only electrophysiological recordings provide the temporal precision required to capture the rich temporal dynamics of neural activity. Movement-related changes in cortical sensorimotor β-activity (13-30 Hz, perimovement^1^ power decrease and post-movement power increase) have been shown to play a key role in movement^3^, making them, putatively, a neural marker for motor learning. However, their role in motor plasticity remains largely unknown.

So far human studies have provided only indirect, and partly contradictory, evidence for behaviourally-relevant learning-induced changes in sensorimotor β-activity^4–7^. While animal studies have identified somatostatin-expressing interneurons in the motor cortex as a key regulator of motor learning^8^, which, in the primary visual cortex, have been related to preferentially drive β-oscillations^9^, direct evidence for changes in β-activity in sensorimotor areas driving motor learning has yet to be established.

Here, we used MEG to directly capture practice-induced functional changes in temporal dynamics during learning of a self-paced, and thus more naturalistic, sequential visuo-motor task (**Supplementary Fig.1, Supplementary Video, Supplementary Methods**, N=19). Investigating human cognition and behaviour under more naturalistic conditions is crucial as it confers greater ecological validity, vital if the results are to be extended to more general principles of motor learning. At the behavioural level, practice leads to faster at constant inter-gate intervals (reduced movement time, *F_5,90_*=7.35, *p*<0.001, 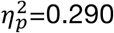, **Fig.1a**) and more accurate (reduced deviation from target, *F_5,90_*=2.89, *p*=0.018, 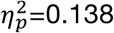, **Fig.1a**) execution of the task.

At the neural level, conventional Fourier-based, wavelet analysis of the first principal component from twelve MEG planar gradiometers pairs covering the sensorimotor cortex (**Supplementary Methods**) provides weak indications that motor learning strengthens the relationship between single-trial time-course of grip force data and β-power (*F_5,90_*=2.11, *p*=0.071, 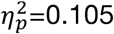, **Supplementary Fig.2a**). Besides, we found practice-induced changes in peri- and post-movement β-power (peri: *F_5,90_*=4.84, *p*<0.001, 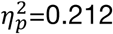; post: *F_5,90_*=2.33, *p*=0.048, 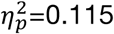; **Fig.1b**), but neither perimovement changes (movement time: *pi_17_*=0.33, *p*=0.382; accuracy: *pi_18_*=0.42, *p*=0.162, **Fig.1c**) nor post-movement changes (movement time: *pi_18_*=-0.27, *p*=0.598; accuracy: *pi_18_*=-0.47, *p*=0.107, **Supplementary Fig.2b**) correlated significantly with behavioural improvement.

**Fig.1.**
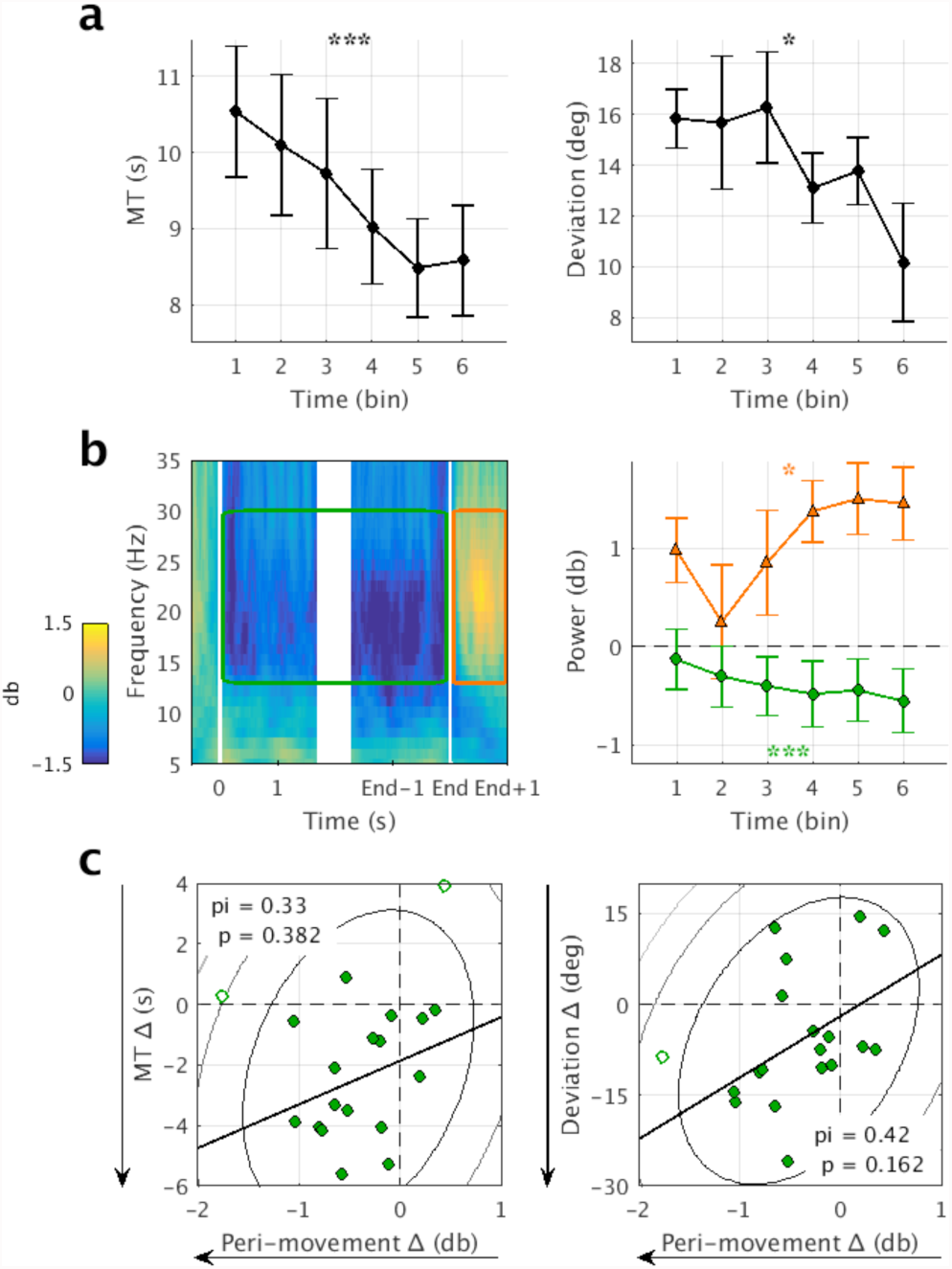
Practice yields improved performance and differential changes in peri- and post-movement Fourier-based β-power. To assess practice-related changes visuomotor task data were divided into six bins. (**a**) Participants improved in terms of decreased movement time (MT, left) and increased accuracy (less deviation, right) with practice. Error bars represent standard error across individuals. (**b**) Group averaged movement-related time-frequency plot (left). As trials are of different length (see Supplementary Fig.3 for intra-and inter-individual differences in movement time) for illustrative purpose data from -0.5 to 1.5 s relative to movement onset and from -1.5 to 1 relative to movement offset are depicted. Two time windows of interest are highlighted: peri-movement (movement onset to movement offset, green) and post-movement (movement offset to 1 s after movement offset, orange). Practice-induced changes (right) in Fourier-based β-power (green circles and orange triangles correspond to peri- and post-movement β-power). Error bars represent standard error across individuals. (**c**) Across-subject brain-behaviour correlations between practice-induced change in peri-movement β-power and practice-induced change in behavioural measures (left: movement time, right: accuracy). Practice-induced change is defined as difference between the last and the first bin. Counter lines represent the bootstrapped Mahalanobis distance from the bivariate mean in steps of six squared units. Open circles represent outliers and the solid line is the regression over the data after outlier removal. Shepherd’s *pi* correlations are reported. Direction of the arrows indicates the expected learning-related change. * indicates a *p*-value < 0.05 and *** a *p*-value < 0.001.

Conventional Fourier-based, wavelet analysis relies on pre-specification of a wavelet length parameter, which constrains the time-scale over which temporal dynamics can evolve. Whilst this provides a good representation of tasks with consistent timing over trials it is likely to be too restrictive for self-paced tasks like the one here, where timings are inherently different across trials. In order to account for this, and hence to better capture practice-induced changes in temporal dynamics (**Supplementary Fig.3a,b**), we further applied state allocation using the Hidden Markov Model (HMM) methodology^10–12^. The HMM estimates state visits whose occurrences and durations are learnt adaptively from the data without pre-specification or constraining the time-scale of dynamics with the data. Specifically, we used univariate autoregressive HMM (HMM-AR) on the first principal component of the group-concatenated single trials from twelve MEG planar gradiometers pairs covering the sensorimotor cortex to describe the data in a probabilistic way using a set of states, where each state is characterized by a distinct spectro-temporal profile (**Supplementary Fig.4** and **Supplementary Methods**).

Following previous work, we inferred three HMM states (blue, red and yellow; see **Supplementary Fig.5** and **Supplementary Discussion** for different numbers of inferred HMM states)^11^. Among the three HMM states, the blue HMM state is characterized by a clear β-power spectral peak (**Fig.2a**, **Supplementary Fig.6**), why hereinafter we refer to it as β-HMM state. The β-HMM state exhibits a distinct movement-related modulation in fractional occupancy (i.e. the across-trial average of the HMM state time-courses, **Supplementary Methods**, **Fig.2b**), and a significant positive correlation with Fourier-based β-power (*t_18_*=17.05, *p*<0.001, one sample t-test, two-sided; **Supplementary Fig.7**). Trial-by-trial correlations revealed that the relationship between β-HMM state fractional occupancy and our behavioural measure of grip force is significantly different from zero (*t_18_*=3.01, *p*=0.008, one sample t-test, two-sided; **Fig.2c**) – this is a significantly stronger correlation than that between Fourier-based β-power and grip force data (*t_18_*=6.42, *p*<0.001, paired samples t-test, two-sided).

**Fig.2.**
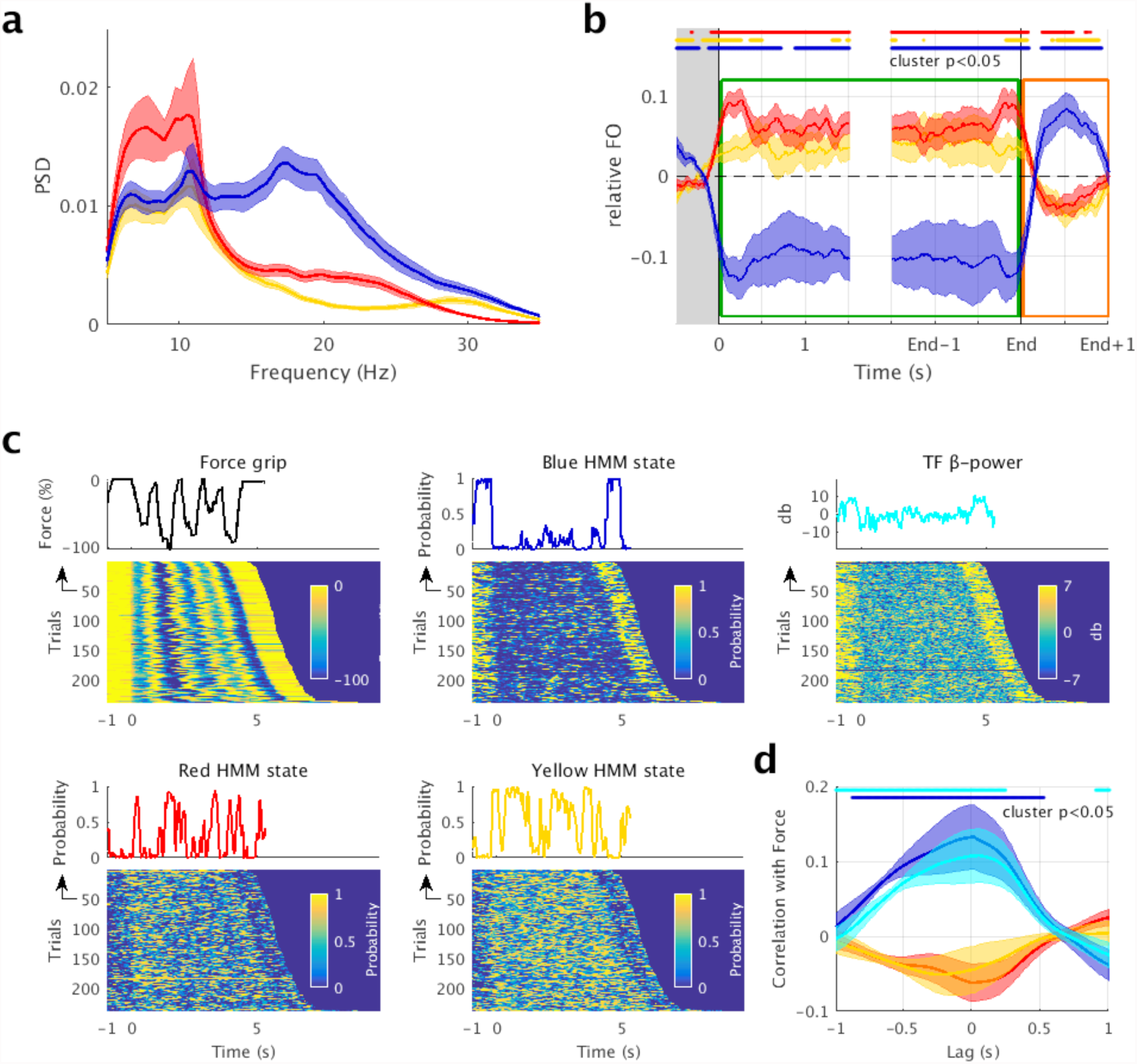
Characterization of the three HMM states. (**a**) Power spectral density (PSD) profiles for each of the three HMM states (blue, red and yellow). Shaded areas represent standard error across individuals. (**b**) Group averaged movement-related fractional occupancy (FO, i.e. the across-trial average of the HMM state time-courses). As trials are of different length (see Supplementary Fig.3 for intra-and inter-individual differences in movement time) for illustrative purpose data from -0.5 to 1.5 s relative to movement onset and from -1.5 to 1 relative to movement offset are depicted. Two time windows of interest are highlighted: peri-movement (movement onset to movement offset, green) and post-movement (movement offset to 1 s after movement offset, orange). Shaded areas represent the standard error across individuals. Significant deviation from zero (*p* < 0.05, one sample t-test, two-sided) is highlighted for each state. Grey shaded area represents the baseline interval. Two time windows of interest are highlighted: peri-movement (green) and post-movement (orange). (**c**) Single-trial time-course of grip force data, time-course of β-power obtained by conventional Fourier-based, wavelet analysis and the three HMM state time-courses qualitatively for one individual. Trials are aligned to movement onset and sorted by movement time. (**d**) Relationship between single-trial time-course of grip force data, time-course of β-power obtained by conventional Fourier-based, wavelet analysis and the three HMM state time-courses quantitatively for the group average. Pearson’s correlations were computed at different lags. Shaded areas represent the standard error across individuals. For the β-HMM state (blue) significant deviation from zero (*p* < 0.05, one sample t-test, two-sided) is highlighted in dark blue and significant deviation from Fourier-based β-power (*p* < 0.05, paired samples t-test, two-sided) is highlighted in light blue.

With practice, the relationship between single-trial time-course of grip force data and β-HMM state fractional occupancy is strengthened significantly (*F_5,90_*=3.46, *p*=0.007, 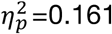; **Fig.3a**), which was not the case of β-power derived from conventional Fourier-based, wavelet analysis. Furthermore, the peri-movement β-HMM state fractional occupancy, lifetime and transitions towards the β-HMM state all decreased; while the post-movement β-HMM state fractional occupancy, lifetime and transitions towards the β-HMM state all increased (**Fig.3b,c,d, Supplementary Fig.9, Table1, Supplementary Results**). Importantly, unlike β-power derived from conventional Fourier-based, wavelet analysis, changes in peri-movement β-HMM state fractional occupancy and changes in performance (i.e., movement time and deviation from target) were related (movement time: *pi_16_*=0.61, *p*=0.026; accuracy: *pi_18_*=0.53, *p*=0.050; **Fig.3e**). There were no brain-behaviour correlations in the post-movement time window (movement time: *pi_17_*=-0.29, *p*=0.510; accuracy: *pi_17_*=-0.17, *p*=1, **Supplementary Fig.2c**)

**Table 1:**
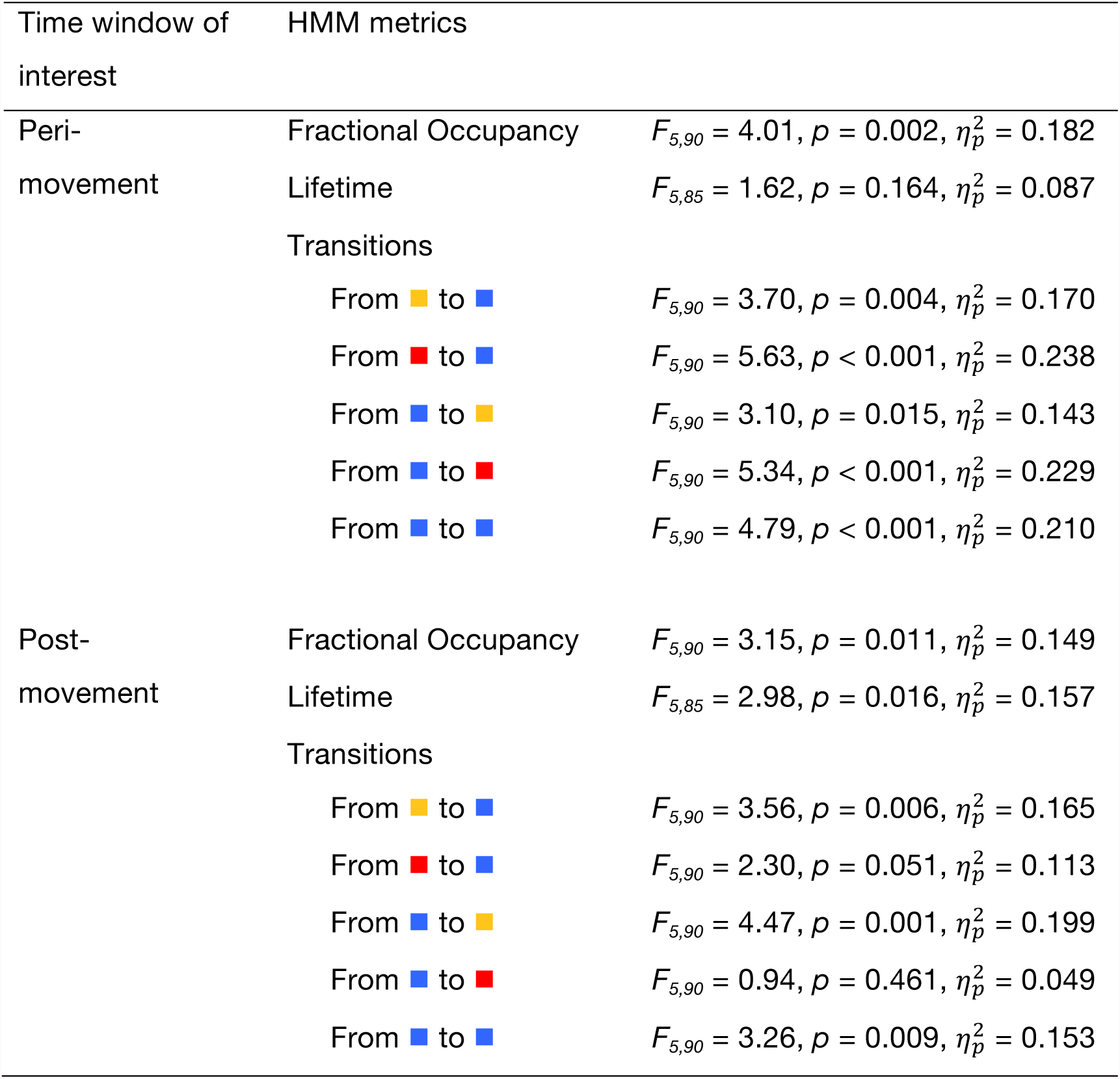
β-HMM state metrics for the two time windows of interest.

.

**Fig.3.**
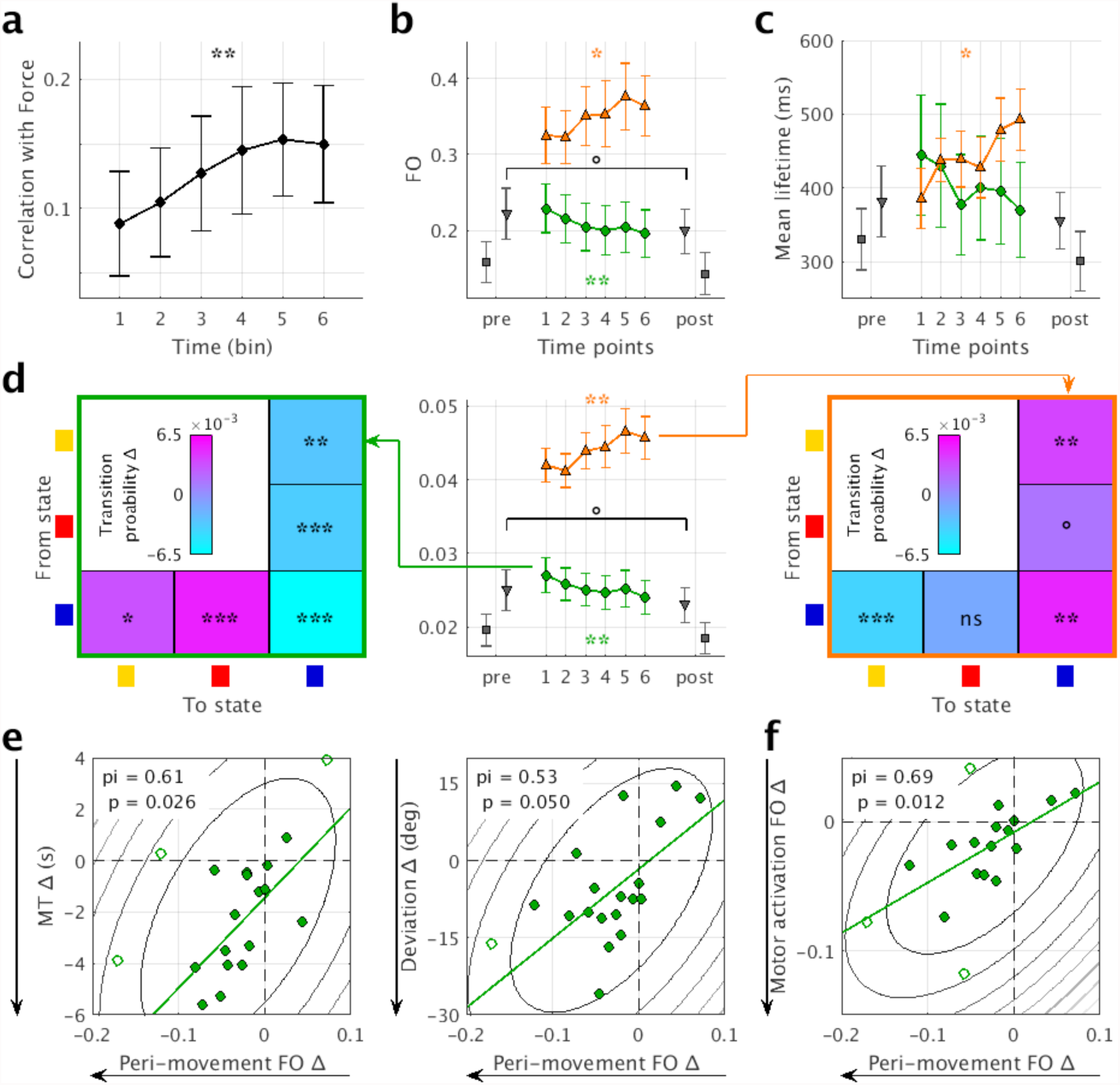
Practice yields differential changes in the β-HMM state (blue in Fig. 2). (**a**) Practice-induced change in the relationship between single-trial time-course of grip force data and β-HMM state fractional occupancy. (**b**) Practice-induced change in fractional occupancy (FO) for the peri-movement (green circles) and post-movement (orange triangles) time window during the visuo-motor task, the simple motor activation task (grey triangles) and resting state (grey squares). Time points refer to pre- and post-learning acquisition of resting state and motor activation task as well as to the six bins of the visuo-motor learning task data. Error bars represent standard error across individuals. (**c**) Practice-induced changes in lifetime. (**d**) Practice-induced changes in peri-movement (green frame) and post-movement (orange frame) transition probabilities. One cell, yellow-HMM state 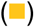 to β-HMM state 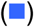, is exemplarily shown in detail. (**e**) Across-subject brain-behaviour correlations between practice-induced change in peri-movement β-HMM state fractional occupancy and practice-induced change in behavioural measures (left: movement time, right: accuracy). Practice-induced change is defined as difference between the last and the first bin. Counter lines represent the bootstrapped Mahalanobis distance from the bivariate mean in steps of six squared units. Open circles represent outliers and the solid line is the regression over the data after outlier removal. Shepherd’s *pi* correlations are reported. Direction of the arrows indicates the expected learning-related change. (**f**) Across-subject brain-behaviour correlations between practice-induced change in peri-movement β-HMM state fractional occupancy and practice-induced change in β-HMM state fractional occupancy during the simple motor activation task. ° indicates a *p*-value < 0.1, * a *p*-value < 0.05, ** a *p*-value < 0.01 and *** a *p*-value < 0.001.

To assess whether the neural changes underpinning learning a visuo-motor task are generalizable to other motor and non-motor contexts, we used the three HMM states descriptions estimated from the visuo-motor task data to investigate dynamics during a simple motor activation task and at rest, both acquired before and after learning (**Supplementary Methods**). Pre-post comparison revealed changes in β-HMM state fractional occupancy (*t_17_*=1.82, *p*=0.086, paired samples t-test, two-sided, grey triangles in **Fig.3b**) and transitions towards the β-HMM state (from yellow HMM state 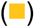 to β-HMM state 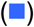: *t_17_*=1.88, *p*=0.077; from red HMM state 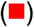 to β-HMM state 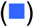: *t_17_*=1.97, *p*=0.065, paired samples t-test, two-sided) during simple motor activation, whereby the direction of the changes were the same as for the peri-movement changes during the visuo-motor task (grey triangles in **Fig.3d, Supplementary Fig.9**). To support the finding that these results generalise across tasks, peri-movement fractional occupancy during motor learning and pre-post learning changes in fractional occupancy during simple motor activation were positively related (*pi_15_*=0.69, *p*=0.012, **Fig.3f**). No similar changes could be observed at rest (all *p-values*>0.1, grey squares in **Fig.3b,c,d** and **Supplementary Fig.9**). No behaviourally-relevant practice-induced changes were observed for the other two HMM states, during the baseline time window, or at a visual control ROI, indicating spectral, temporal and spatial specificity in the observed findings (**Supplementary Fig.10,11,12** and **Supplementary Results**).

We have found strong evidence that motor practice induces differential changes in the temporal dynamics of cortical sensorimotor β-activity (as captured by the HMM metrics), indicating behaviourally-relevant alterations in the activity of local intracortical microcircuits. In general, GABAergic interneurons have been suggested as major players in generating and regulating neural oscillations, whereby the subtype somatostatin-expressing interneuron has been specifically related to orchestrating β-oscillations^9^. Both peri- and post-movement movement-related changes in the temporal dynamics of cortical sensorimotor β-activity were found to be significantly more pronounced over the course of practice; consistent with the general theory that, during the initial stage of learning, additional resources are required as the skill is not yet automated^13^. Brain-behaviour correlation analyses suggest that peri-movement changes, but not post-movement changes, are behaviourally relevant for the learning of this sequential movement (**Supplementary Discussion**). Thus, our findings demonstrate a direct role of peri-movement changes in the temporal dynamics of cortical sensorimotor β-activity in motor learning, suggesting a functional role of β-activity beyond maintaining the current sensorimotor set, or “status quo”^14^.

One of the most important questions surrounding any demonstration of the neural basis of plasticity is whether it can be generalised to other motor and non-motor contexts. Here, we demonstrate that the behaviourally-relevant neural signature of motor learning showed positive transfer to a simple movement task, but not to rest. This state-selectivity is consistent with the long-held principle of identical elements^15^, which states that the level of transfer depends, among other factors, on the similarity between training and transfer contexts.

Here, we have used HMM metrics to provide a better reflection of changes in movement-related temporal dynamics than β-power derived from conventional Fourier-based, wavelet analysis. There are several non-mutually exclusive reasons to support the implementation of this approach, among them that the HMM delivers a better signal-to-noise ratio and more spectral features. Additionally, these results may be relevant for the incipient debate on the interpretation of frequency-specific patterns of neural activity; the superiority of HMM speaks rather for the existence of transient bursts or a hybrid combination of bursts and sustained rhythms than for pure rhythmically sustained oscillations^16^, though this hypothesis remains to be directly tested.

Cellular and mechanistic interpretations of the observed changes are vital for translational propositions, but are difficult given the current state of research. Findings from animal studies suggest that Hebbian long-term potentiation, through inhibition of distal dendrites by somatostatin-expressing inhibitory neurons, is one possible mechanism for learning-related changes in the temporal dynamics of β-activity^8,9,17,18^. Our results underscore the importance of sensorimotor β-activity in motor plasticity, but are not meant to imply that activity in other brain areas (e.g. cortico-striatal and cortico-cerebellar system) are not altered in the course of motor learning. Parallel human and animal studies covering micro- to macroscopic scales and integrating *in vitro* and *in vivo* together with computational modelling are required to determine the neurobiological underpinnings of practice-induced plasticity in the temporal dynamics of neural activity.

## Acknowledgments

We thank Anna C Nobre for helpful discussions and suggestions on behavioural data analysis. This research was funded by a Sir Henry Dale Fellowship to CJS, funded by the Wellcome Trust and the Royal Society (102584/Z/13/Z), and supported by the NIHR Oxford Health Biomedical Research Centre, and the John Fell Fund. The Wellcome Centre for Integrative Neuroimaging is supported by core funding from the Wellcome Trust (203139/Z/16/Z). MWW is supported by the Wellcome Trust (106183/Z/14/Z) and the MRC UK MEG Partnership Grant (MR/K005464/1). DV is supported by a Wellcome Trust Strategic Award (098369/Z/12/Z). JS was supported by a Wellcome Trust, Four-Year PhD Studentship (092759/Z/10/Z).

## Author contributions

CJS and JS designed the study. ELH, LJ, CJS and SB collected the data. CZ analysed the data. CJS supervised the project. MWW, RB, AJQ and DV provided assistance with data analysis and interpretation. CZ wrote the manuscript and all of the authors edited the manuscript.

## Supplementary methods

**Supplementary Fig.1:**
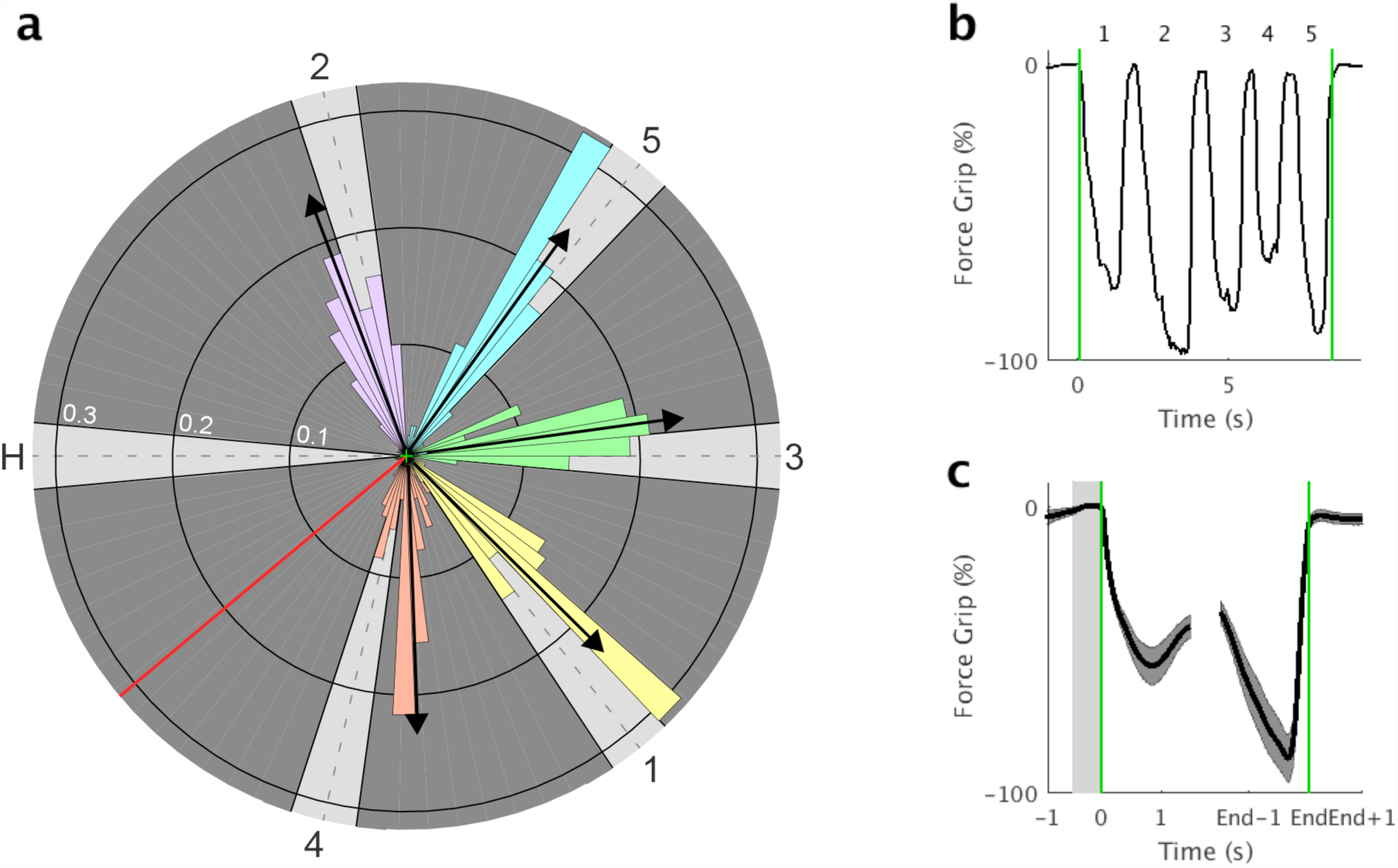
Design of the visuo-motor task overlaid with distribution of accuracy. Participants saw the dark grey circle with its light grey numbered gates and a red cursor over the whole time period of the visuo-motor task. Rotation of the on-screen cursor (red line) was controlled by squeezing (anti-clockwise) and relaxing (clockwise) a force transducer with the right hand (Supplementary Video). The goal of the task was to move the cursor quickly and accurately between the start/end position (Home) and a numbered order of gates (Home-1-Home-2-Home-3-Home4-Home-5-Home). Here, the visual display shown to the participants is overlaid with the accuracy histograms of the circular distance between the reversal point of the cursor and the centre of the approached gate. The direction of the arrow indicates the average accuracy for each gate. Note that the distance to the centre of the circle and the centre of the gates are not displayed to the participants. (**b**) Single trial grip force data epoched from 1 s before movement onset to 1 s after movement offset. Green lines indicate movement onset and offset. (**c**) Group averaged grip force data. As trials are of different length (see Supplementary Fig.3 for intra-and inter-individual differences in movement time) for illustrative purpose data from -0.5 to 1.5 s relative to movement onset and from -1.5 to 1 relative to movement offset are depicted. Grey rectangle indicates the baseline interval and shaded areas represent standard error across individuals.

**Supplementary Fig.2:**
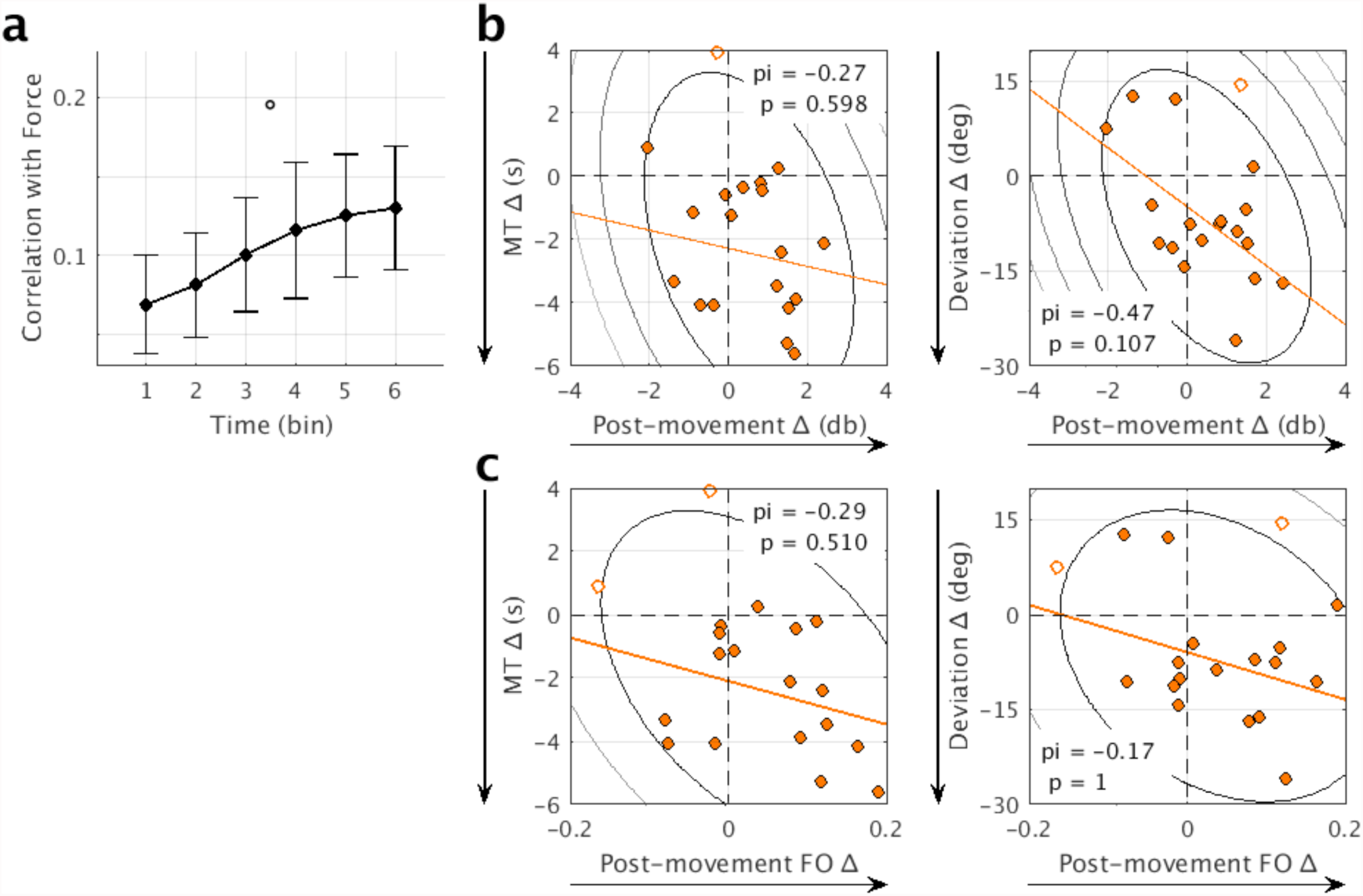
(**a**) Practice-induced change in the relationship between single-trial time-course of grip force data and β-HMM state fractional occupancy. ° indicates a *p*-value < 0.1. (**b**) Across-subject brain-behaviour correlations between practice-induced change in post-movement β-Power and practice-induced change in behavioural measures (left: movement time, right: accuracy). Practice-induced change is defined as difference between the last and the first bin. Counter lines represent the bootstrapped Mahalanobis distance from the bivariate mean in steps of six squared units. Open circles represent outliers and the solid line is the regression over the data after outlier removal. Shepherd’s *pi* correlations are reported. Direction of the arrows indicates the expected learning-related change. (**c**) As b, but for post-movement β-HMM state fractional occupancy.

**Supplementary Fig.3:**
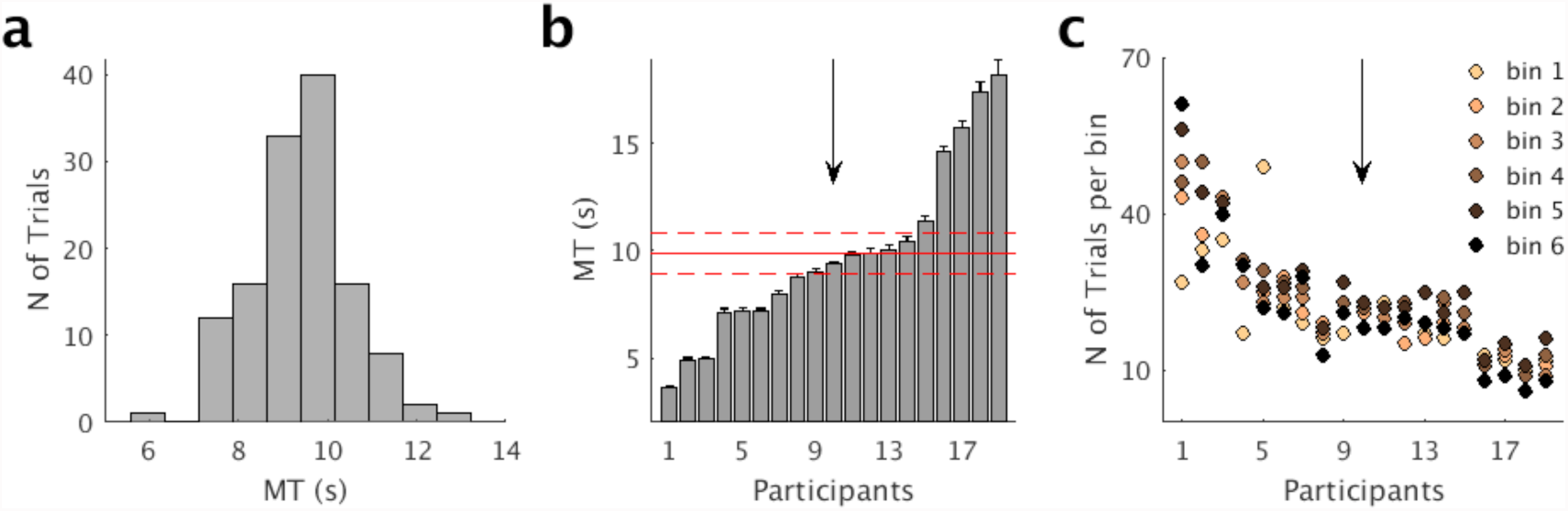
Intra- and inter-individual differences in movement time and number of trials. (**a**) Distribution of movement time (MT) for one representative individual. (**b**) Mean movement time per individual with error bars reflecting standard error. Individuals are sorted based on mean movement time. Horizontal red lines indicate group average plus/ minus one standard error. Arrow indicates the individual displayed in a. (**c**) Number of trials per bin for each participant separately (Supplementary Methods for details). Individuals are sorted based on mean movement time. Arrow indicates the individual displayed in (a).

**Supplementary Fig.4:**
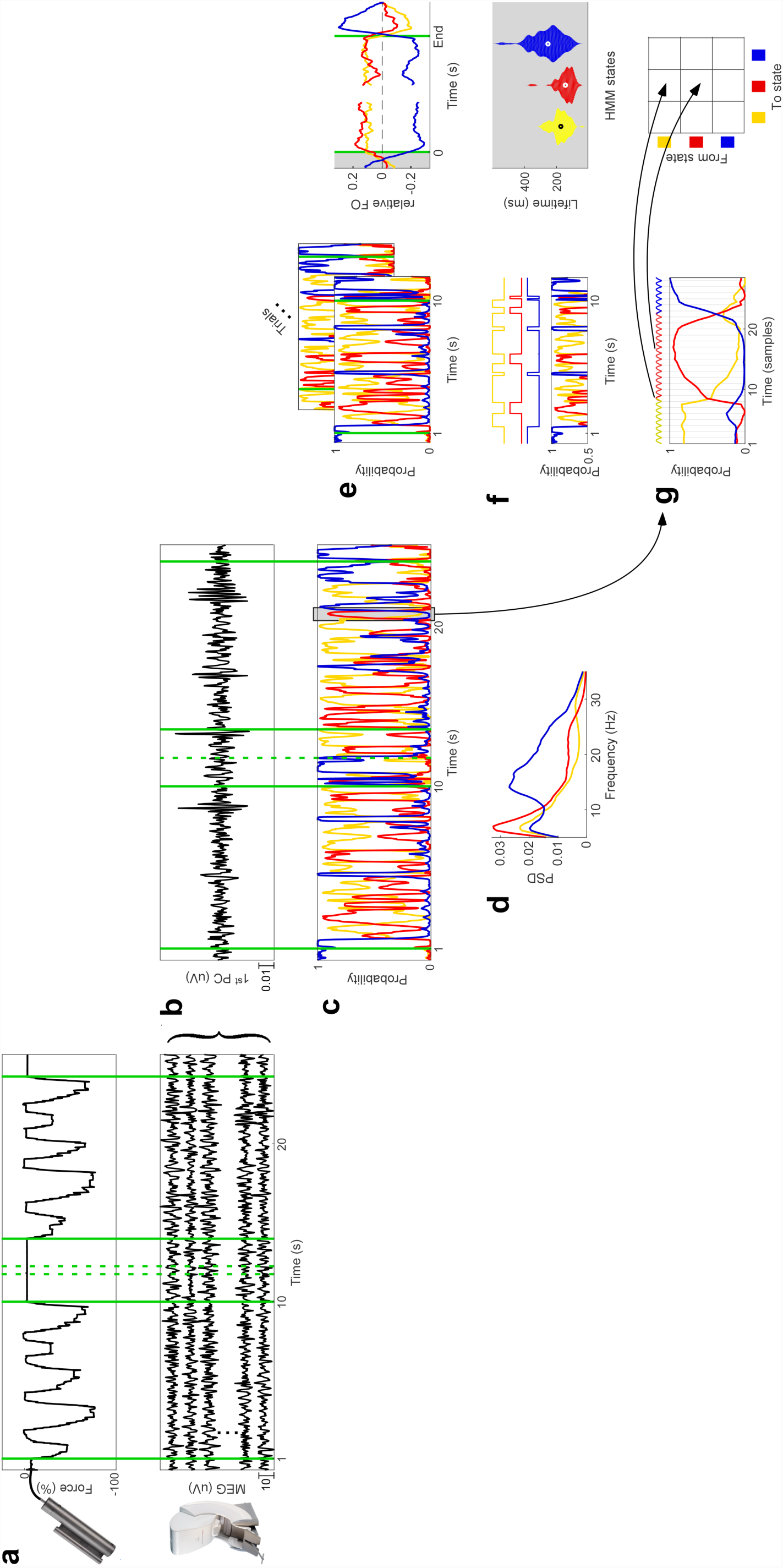
From raw data to HMM metrics: graphical representation of HMM analysis pipeline for the visuo-motor task data. (**a**) MEG and grip force data were recorded simultaneously and grip force data were used to determine movement onset and offset. (**b**) Following pre-processing (Supplementary Methods for details) the 1^st^ PC of the group-concatenated single trials of the visuo-motor task data from twelve MEG planar gradiometers pairs covering the sensorimotor cortex were submitted to a single-channel autoregressive Hidden Markov Model analysis (HMM-AR). We inferred three HMM states (blue, red and yellow). The HMM-AR provides a time-course (**c**) and a spectrum (**d**) for each of the inferred states. Based on these the following HMM metrics were estimated. (**e**) Average across single trial state time-courses (left) results in fractional occupancy (FO), i.e. across-trial average of the fraction of time spent in each state (right). (**f**) Lifetimes, computed following Viterbi decoding (left), are define as the trial-by-trial average amount of time spent in each state before transitioning out of that state (right). (**g**) Transition probabilities describe the mean of the trial-by-trial likelihood of transitioning from a given state to the same or another state.

**Supplementary Fig.5:**
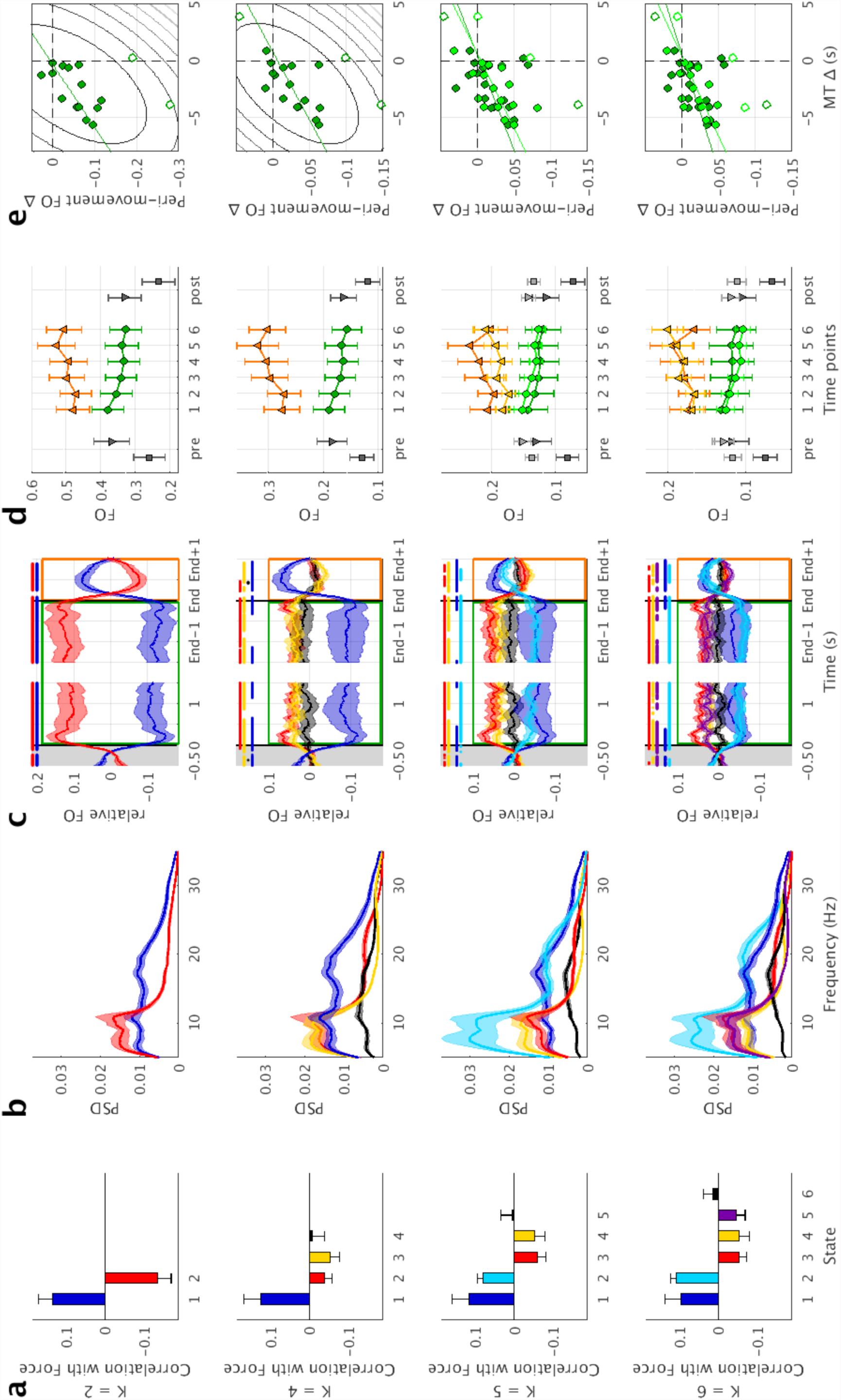
Results are robust across different number of inferred states (K = 2, 4, 5, 6). (**a**) Group averaged Pearson’s correlation between single-trial time-course and time-course of grip force data at lag zero. (**b**) Group averaged spectral profiles of the states. Shaded areas represent standard error across individuals. (**c**) Group averaged movement-related fractional occupancy (FO, i.e. the across-trial average of the HMM state time-courses). Shaded areas represent standard error across individuals. Significant deviation from zero (*p* < 0.05, one sample t-test, two-sided) is highlighted for each state. Grey shaded area represents the baseline interval. Two time windows of interest are highlighted: peri-movement (green) and post-movement (orange). (**d**) Practice-induced change in fractional occupancy for the β-MM state(s). Green circles and orange triangles correspond to peri- and post-movement fractional occupancy during the visuo-motor task. Grey triangles and squares correspond to the factional occupancy during the motor activation task and resting state, respectively. Time points refer to pre- and post-learning acquisition of resting state and motor activation task as well as to the six bins of the visuo-motor learning task data. Dark and light colours correspond to the β-HMM state(s) illustrated in a-c in dark and light blue, respectively. Error bars represent standard error across individuals. (**e**) Across-subject brain-behaviour correlations between practice-induced change in movement time and practice-induced change in peri-movement β-HMM state(s) fractional occupancy during the visuo-motor task. Practice-induced change is defined as difference between the last and the first bin. Counter lines represent the bootstrapped Mahalanobis distance from the bivariate mean in steps of six squared units. Open circles represent outliers and the solid line is the regression over the data after outlier removal.

**Supplementary Fig.6:**
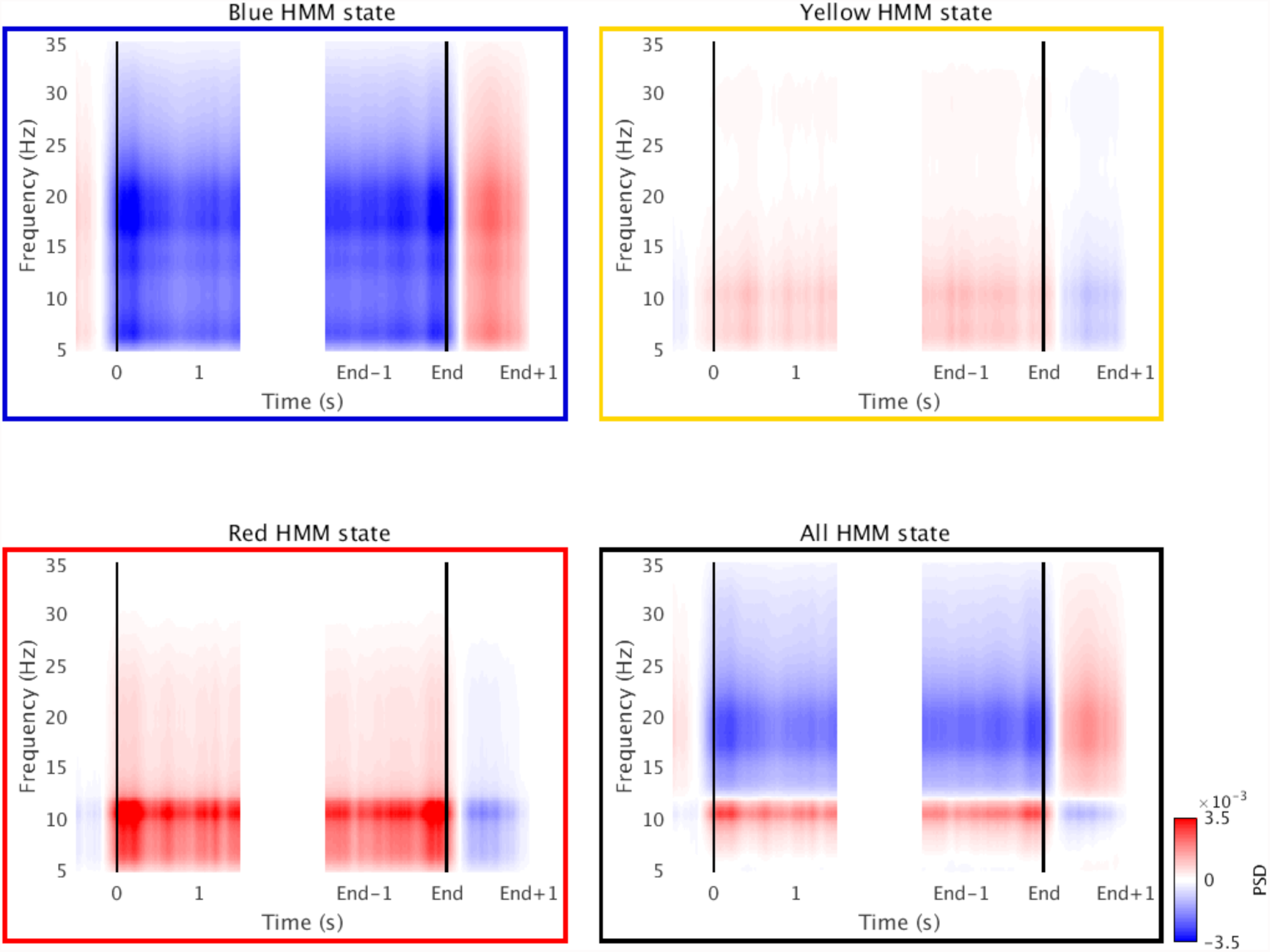
Group averaged movement-related reconstructed HMM-AR regularised time-frequency plots for each of the thee HMM states separately and all HMM states together. As trials are of different length (see Supplementary Fig.3 for intra-and inter-individual differences in movement time) for illustrative purpose data from -0.5 to 1.5 s relative to movement onset and from -1.5 to 1 relative to movement offset are depicted.

**Supplementary Fig.7:**
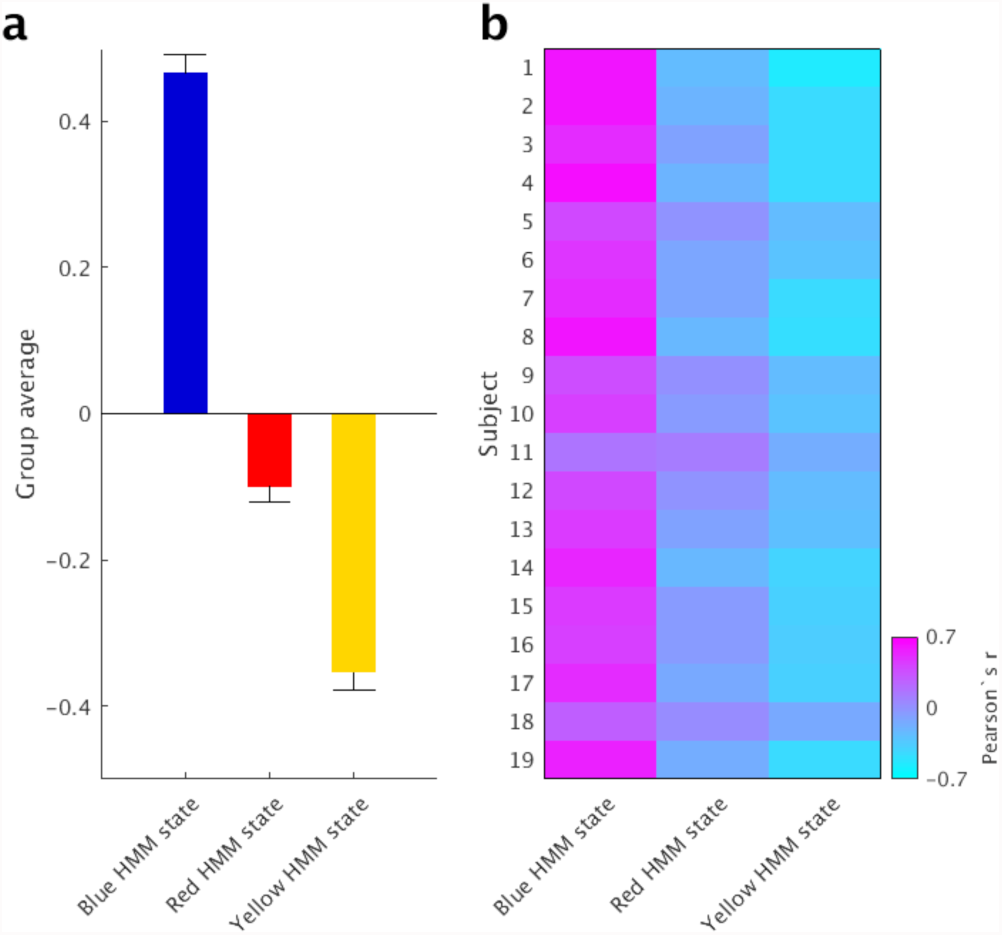
Relationship between the three HMM states and β-power obtained by Fourier-based, wavelet analysis. Group averaged (**a**) and individual (**b**) Pearson’s correlation between single-trial HMM state time-course and time-course of β-power obtained by Fourier-based, wavelet analysis at lag zero. Error bars in represent standard error across individuals.

**Supplementary Fig.8:**
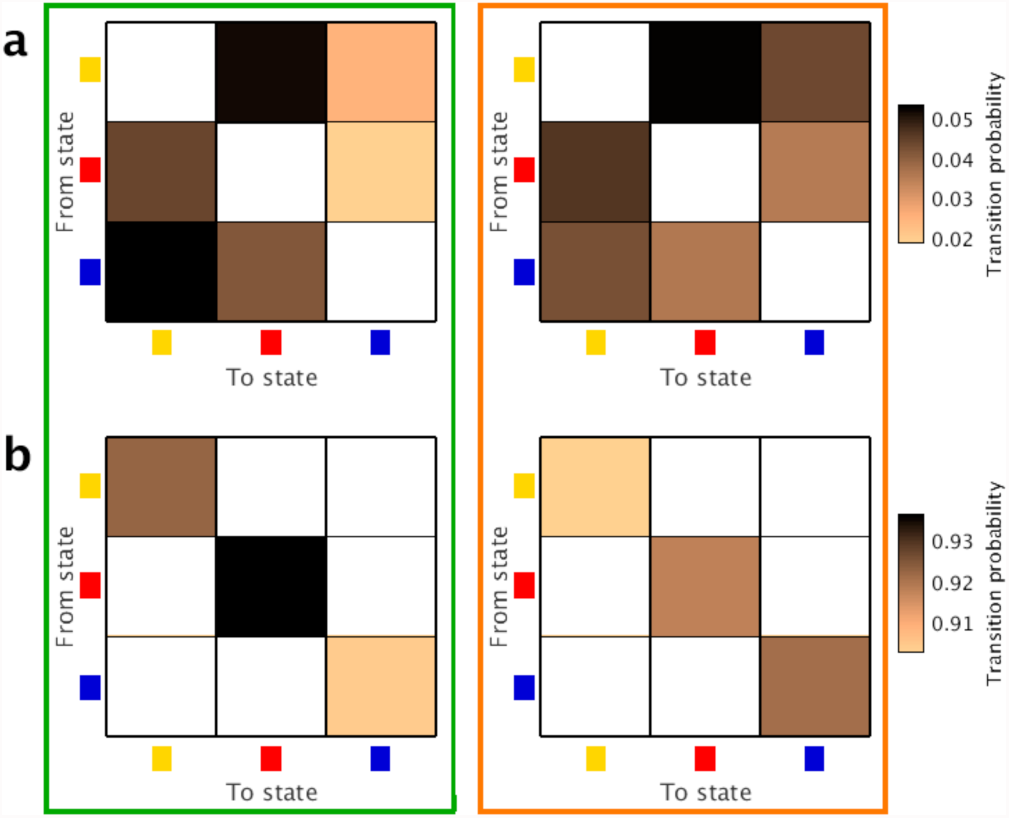
Group averaged transition probability matrix for perimovement (left) and post-movement (right) indicating the probability of transitioning from any state to any state. Note – the matrix is scaled differently for the off-diagonal cells (**a**) and diagonal cells (**b**).

**Supplementary Fig.9:**
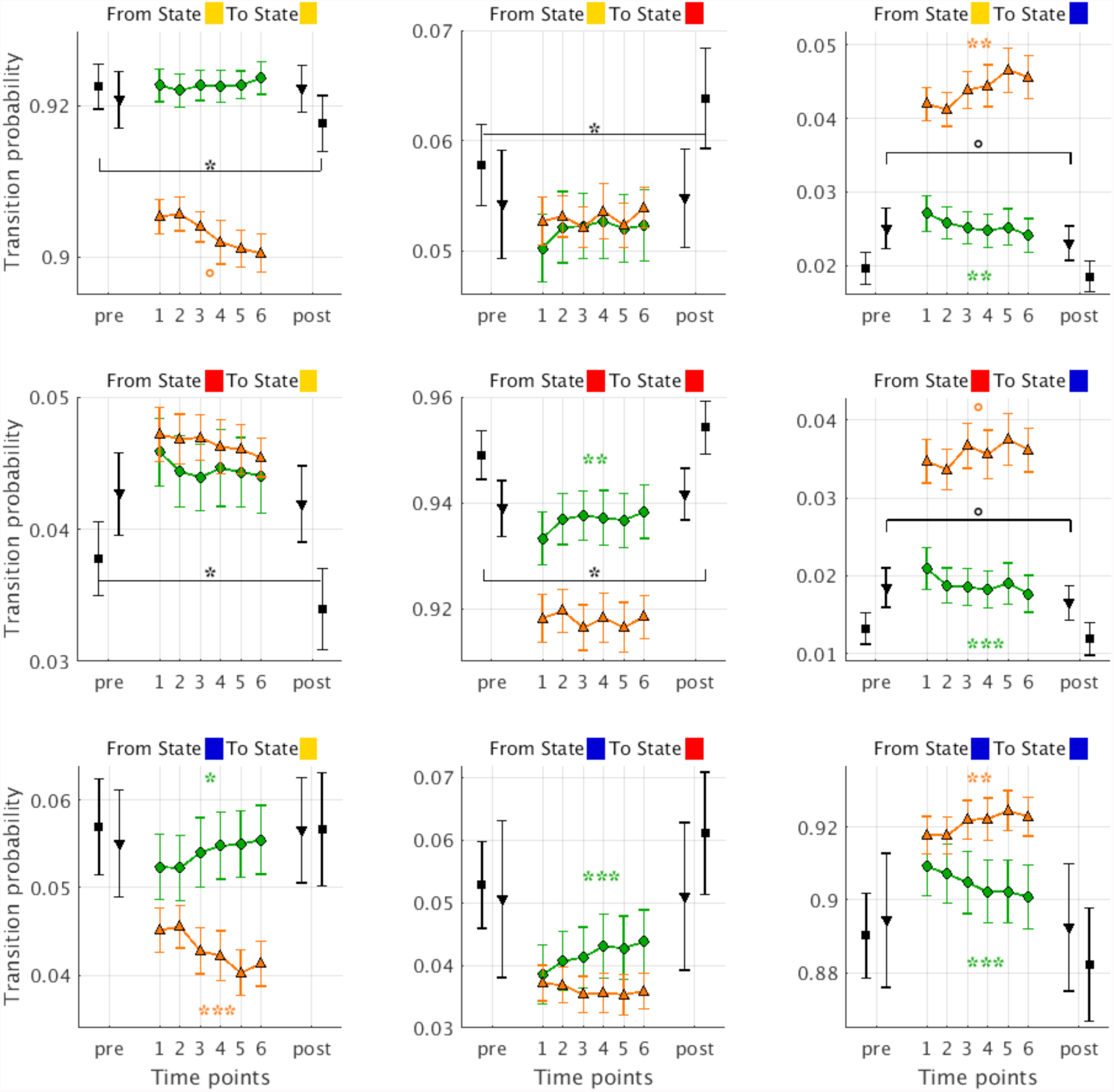
Practice yields differential changes in transition probabilities. Green circles and orange triangles correspond to peri- and post-movement transition probabilities during the visuo-motor task. Grey triangles and squares correspond to the transition probabilities during the motor activation task and resting state, respectively. Time points refer to pre- and post-learning acquisition of resting state and motor activation task as well as to the six bins of the visuo-motor learning task data. Error bars represent standard error across individuals. ° indicates a *p*-value < 0.1, * a *p*-value < 0.05, ** a *p*-value < 0.01 and *** a *p*-value < 0.001.

**Supplementary Fig.10:**
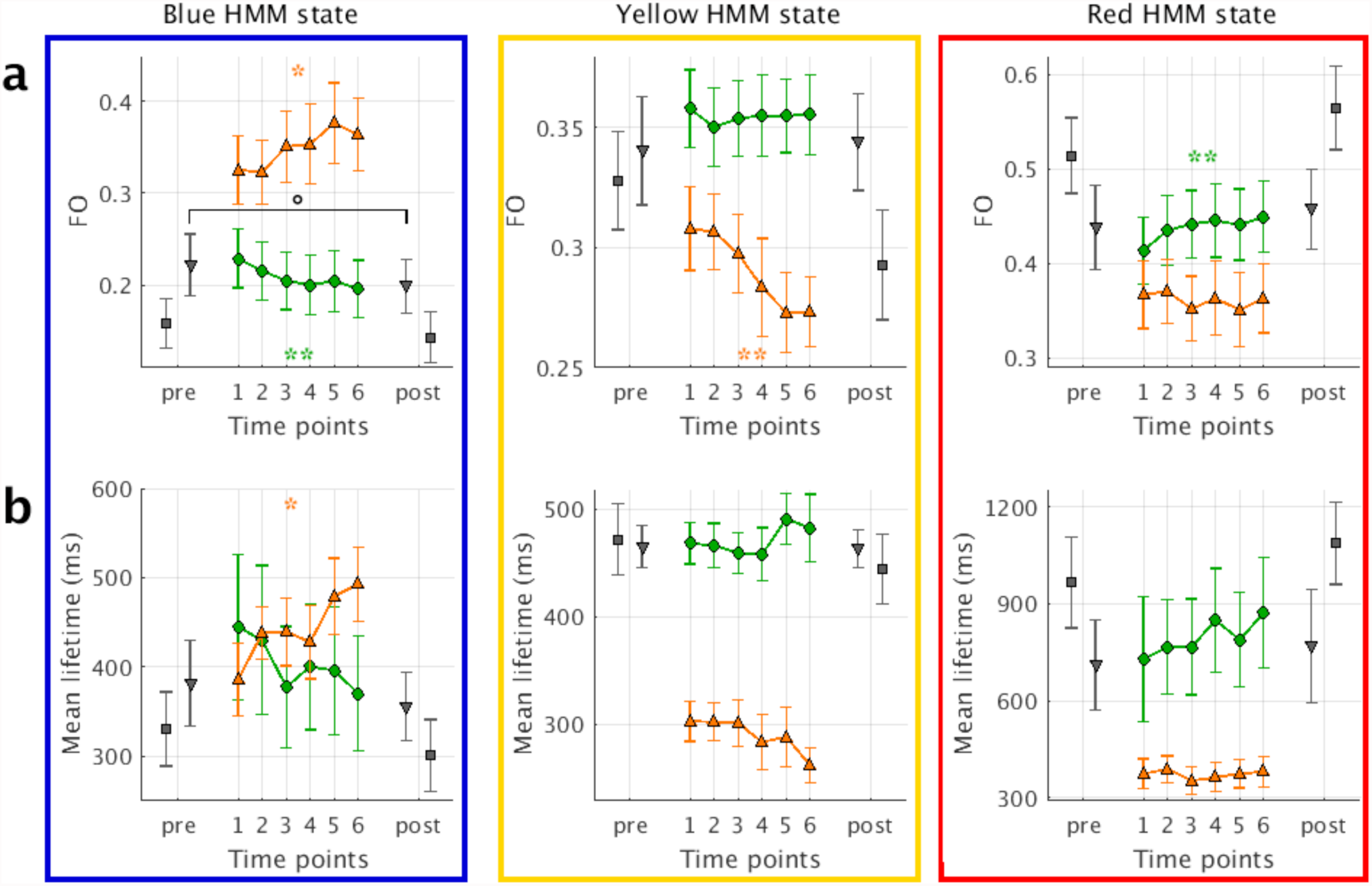
Practice-induced changes in (**a**) fractional occupancy (FO) (**b**) and lifetime for the yellow and red HMM state. In addition, the blue HMM state is depicted for reference. Green circles and orange triangles correspond to peri- and post-movement fractional occupancy/ lifetime during the visuo-motor task. Grey triangles and squares correspond to fractional occupancy/ lifetime during the motor activation task and resting state, respectively. Time points refer to pre- and post-learning acquisition of resting state and motor activation task as well as to the six bins of the visuo-motor learning task data. Error bars represent standard error across individuals. ° indicates a *p*-value < 0.1, * a *p*-value < 0.05 and ** a *p*-value < 0.01.

**Supplementary Fig.11:**
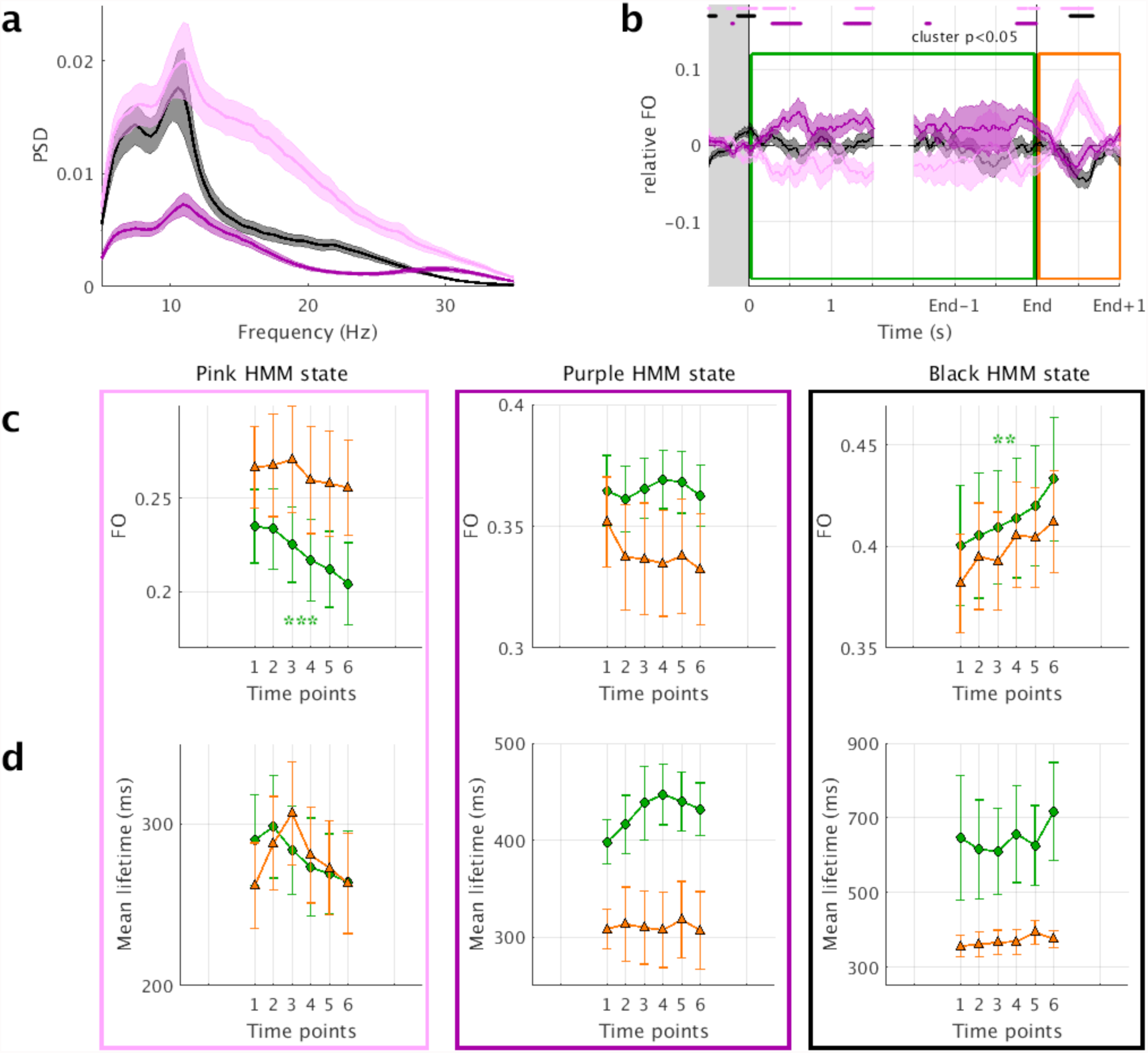
HMM metrics are estimated from a visual control ROI using three HMM states derived from the same visual ROI. (**a**) Group averaged spectral profiles of the states. Shaded areas represent the standard error across individuals. (**b**) Group averaged movement-related fractional occupancy (FO, i.e. the across-trial average of the HMM state time-courses). As trials are of different length (see Supplementary Fig.3 for intra -and inter-individual differences in movement time) for illustrative purpose data from -0.5 to 1.5 s relative to movement onset and from -1.5 to 1 relative to movement offset are depicted. Shaded areas represent standard error across individuals. Significant deviation from zero (*p* < 0.05, one sample t-test, two-sided) is highlighted for each state. Grey shaded area represents the baseline interval. Two time windows of interest are highlighted: peri-movement (green) and post-movement (orange). Changes in fractional occupancy (**c**) and lifetime (**d**). Green circles and orange triangles correspond to peri- and post-movement fractional occupancy/ lifetime during the visuo-motor task. Time points refer to pre- and post-learning acquisition of resting state and motor activation task as well as to the six bins of the visuo-motor learning task data. Error bars represent standard error across individuals. ** indicates a *p*-value < 0.01 and *** a *p*-value < 0.001.

**Supplementary Fig.12:**
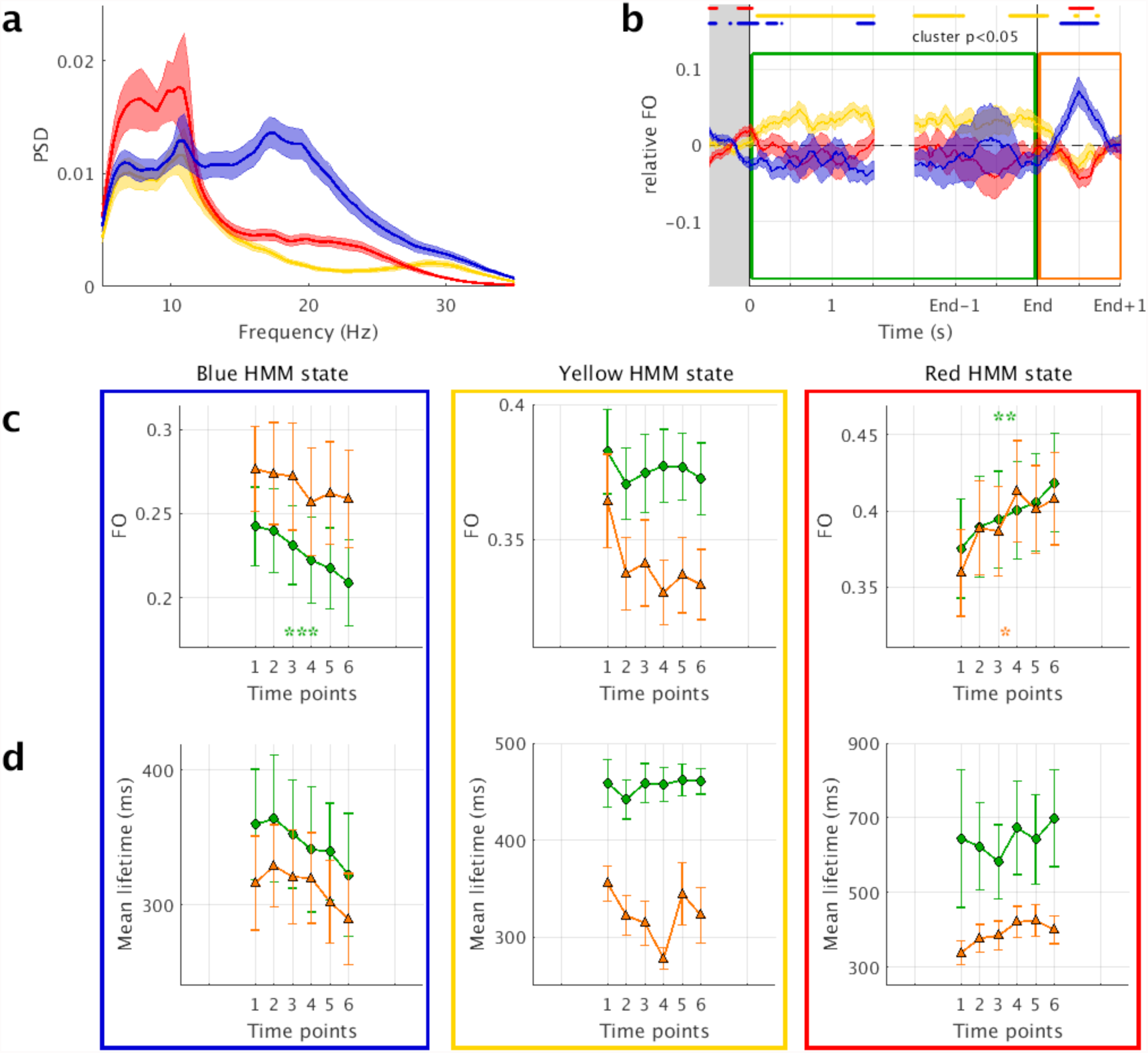
As Supplementary Fig. 11, HMM metrics are estimated from a visual control ROI, but this time using the three HMM states derived from the sensorimotor ROI. * indicates a *p*-value < 0.05, ** a *p*-value < 0.01 and *** a *p*-value < 0.001.

**Supplementary Fig.13:**
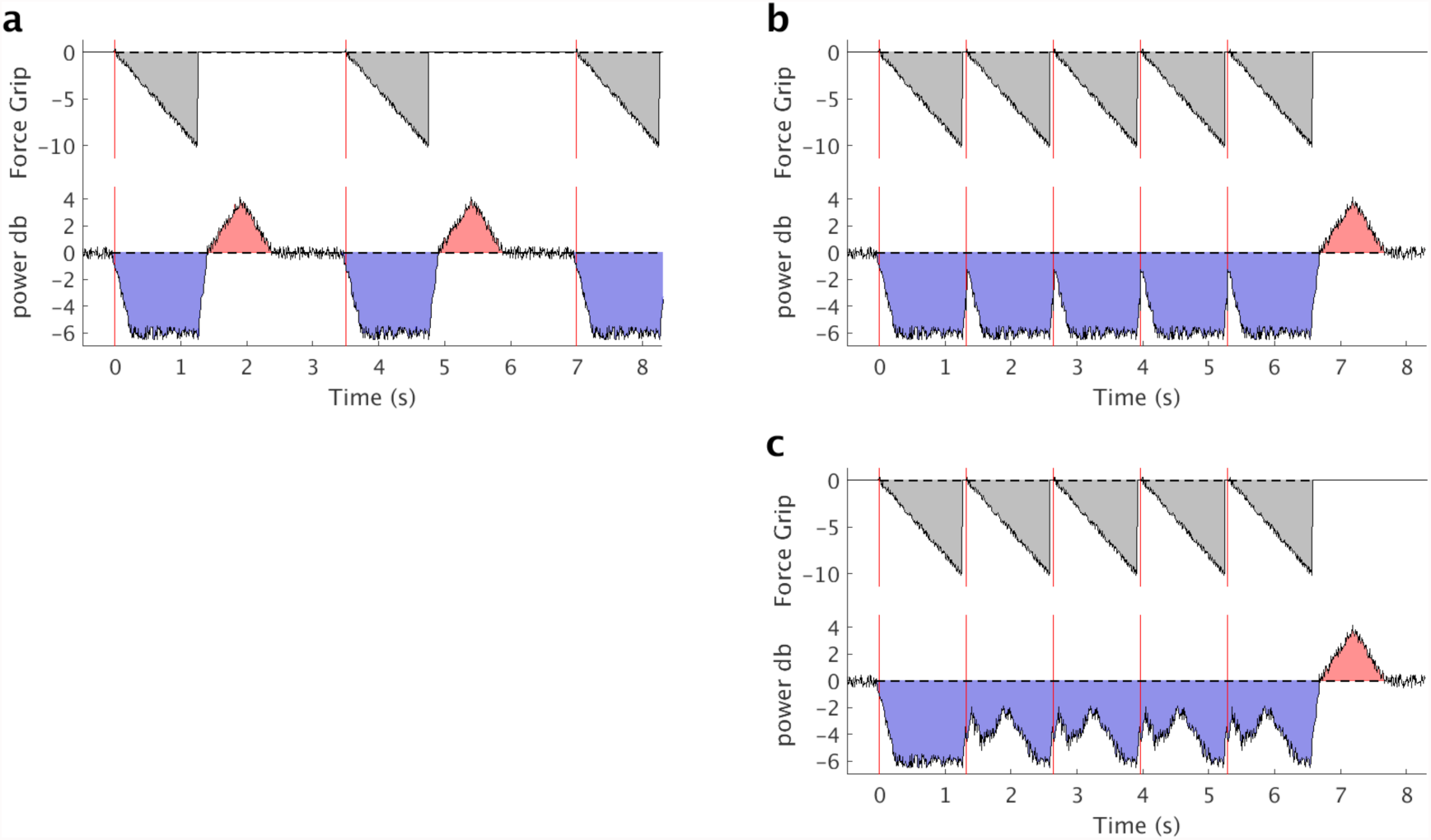
Schematic illustration of possible movement-related changes in temporal dynamics of cortical sensorimotor β-activity for sequential movements. (**a**) Sequential movement with long inter-movement intervals. Perimovement decrease and post-movement increase in power are highlighted in blue and red respectively. Red vertical lines indicate movement onsets. (**b**) Sequential movement with short inter-movement intervals with interruption of post-movement increase in power by the pre- and peri-movement decrease in power of the subsequent movement. (**c**) Sequential movement with short inter-movement intervals with cancelation of post-movement increase in power and pre- and perimovement decrease in power of the subsequent movement.

### Participants

19 healthy volunteers (age range 18-31 years; mean age 23.67 years, standard deviation 4.20 years; 9 females) participated in this study, which was approved by the Central Oxfordshire Research Ethics Committee (MSD-IDREC-C1-2014-053), and in accordance with the Declaration of Helsinki. Informed written consent was obtained from all participants. All participants were free of neurological and psychiatric disorders, right-handed, as determined by the Edinburgh Handedness Questionnaire and had normal or corrected-to-normal vision.

### Experimental Design

Each session started with questionnaires followed by MEG data acquisition. MEG data comprised 8.5 min. pre-learning resting state, 5 min. pre-learning motor activation task, 24 min. visuo-motor learning, 8.5 min. post-learning resting state and 5 min. post-learning motor activation task.

### Motor Learning

All participants underwent 24 min. of repeated motor task practice. Throughout this time a dark grey circle with six light-grey 10° wide segments (Home: 175-185°, Gate 1: 304-314°, Gate 2: 98-108°, Gate 3: 355-5°, Gate 4: 252-262° Gate 5: 46-56°) and a red cursor (length = *r*) were constantly displayed to the participant (**Supplementary Fig.1, Supplementary Video**). Stimuli were created and presented using Neurobehavioral Systems Presentation software. Participants held a force transducer (Current Designs, Philadelphia, USA) in their right hand, which was rested on a pillow on their lap. Squeezing the force transducer moved the curser anti-clockwise, while relaxing caused the curser to move clockwise. The goal of the task was to move the cursor quickly and accurately between the start/end position (Home) and a sequential order of gates (Home-1-Home-2-Home-3-Home-4-Home5-Home) by modulating the force exerted onto the transducer. Movement time (MT; time from movement onset, i.e. initiation to visit Gate 1, to movement offset, i.e. arrival at Home following gate 5 visit) and accuracy (i.e. absolute angular difference between the centre of the gate and the reversal point of the cursor) were extracted as behavioural measures.

Aiming to facilitate learning, both, time point of trial initiation and trial duration were completely self-paced, the sole exception being an imposed inter-trial-interval of at least 1 s. While this made the task appearing very natural, it also means that given a similar total movement time (MT) across participants 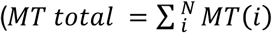, where N denotes the number of trials per individual, *M* = 19.70 min, *SE* = 24.88 s), but substantial inter- and intra-individual differences in movement time (**Supplementary Fig.3a,b**), the number of completed trials differs across participants.

To assess practice-induced changes data were divided into six bins of equal movement time (i.e., 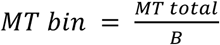, where B denotes the number of bins, *M* = 3.28 min, *SE* = 4.15 s, **Supplementary Fig.3c**), and mean movement time as well as mean accuracy compared using repeated measures analysis of variance (ANOVA) with bin as within-subjects factor.

### Motor activation task

The total duration of the motor activation task was 8.5 min. A phase-encoding task was used, which involved continuous button presses of individual digits of the right hand (D2: index, D3: middle, D4: ring, D5: little) on an MEG-compatible button box (Current Designs, Philadelphia, USA) with no rest periods. Participants were instructed to press each digit eight times, at a rate of 1 Hz. The forward version of the task cycled from D2 to D5 inclusive, with the resulting 32 cycle repeated eight times. The backward version of the task was identical in duration but cycled from D5 to D2 inclusive. The two versions were pseudorandomized across participants. Participants were presented with four white circles, corresponding to the four digits. The displayed circles flashed individually at a constant frequency to cue participants to make button presses at the specified rate. Stimuli were created and presented using Neurobehavioral Systems Presentation software. The paradigm has been applied and validated using fMRI previously^1,2^

### Resting state

Participants were asked to sit still in the MEG scanner with their eyes fixating a cross.

### MEG data acquisition

Whole-head MEG recordings were acquired in a magnetically shielded room using a 306-channel VectorView System (Elekta Neromag, Elekta, Stockholm, Sweden), with one magnetometer and two orthogonal planar gradiometers at each of 102 locations, at Oxford Centre for Human Brain Activity. Head position was monitored at the beginning of each acquisition type using four head position indicator (HPI) coils attached to the scalp. HPI coil locations, head positions from across the scalp and three anatomical fiducial locations (nasion, left and right pre-auricular points) were digitalized using a magnetic Polhemus FastTrak 3D system (VT, USA) before data acquisition. Vertical and horizontal electo-oculogram as well as electrocardiogram were also measured to detect eye blinks, horizontal eye movements and heartbeat, respectively. MEG data were sampled at 1000 Hz using a 0.03-330 Hz band-pass filter during digitalisation. Stimuli were back projected (Panasonic PT D7700E, Panasonic, Osaka, Japan) on a 43 × 54.5 cm screen placed 120 cm in front of the participant, with a spatial resolution of 1280 × 1024 and a refresh rate of 60 Hz.

### MEG data analysis

MEG data acquired during visuo-motor learning, pre- and post-resting state and pre- and post-motor activation were pre-processed separately. External noise was reduced from MEG data by means of spatio-temporal signal-space separation (TSSS) and head movements (detected using HPI coils) corrected, both using MaxMove software as implemented in MaxFilter version 2.1 (Elekta Neromag, Elekta, Stockholm, Sweden). Further MEG data analyses were performed using the in-house OHBA Software Library (OSL: https://ohba-analysis.github.io/osl-docs/) version 2.2.0. Continuous data were down-sampled to 250 Hz to reduce computational demands. To optimize independent component analysis (ICA) decomposition quality we performed several preprocessing steps before applying ICA. First, slow trends, high frequencies and line noise were removed by applying a band-pass (0.1-100 Hz) and a notch (48-52 Hz) filter. Second, data containing non-stereotypical artefacts were removed. This was achieved by epoching the data into continuous non-overlapping 1 s intervals and rejecting those containing non-stereotypical artefacts using a generalized extreme Stundentized deviate method^3^ at a significance level of 0.05 with a maximum number of outliers limited to 20% of the data. This processing is considered an optimal trade-off for retaining much variance in the data but preventing the ICA algorithm being adversely affected by low- and high-frequency contributions as well as gross artefacts. Remaining data were used to estimate the unmixing weights of 64 independent components using temporal FastICA across sensors^4^. Independent components representing stereotypical artifacts such as eye blinks, eye movements, and electrical heartbeat activity were manually identified and regressed out of the maxfiltered data. Second, cleaned maxfiltered data were down-sampled to 100 Hz, band-pass filtered (5-35 Hz). Data acquired during the visuo-motor learning task were epoched with respect to grip force data (from 1 s before movement onset to 1 s after movement offset; see **Supplementary Fig.1b,c**), while data acquired during resting state and the motor activation task were kept continuous.

Fourier-based, wavelet analysis and hidden markov modelling were performed on the first principal component of an *a priori* anatomically defined ROI comprising twelve planar gradiometer pairs covering left sensorimotor cortex (explained variance: *M* = 33.16; *SD* = 2.84). In order to test the spatial specificity of the effects a control ROI, comprising ten planar gradiometer pairs above the occipital pole served as control region (explained variance: *M* = 37.20; *SD* = 10.30), was defined.

Time-frequency decomposition of the visuo-motor learning data was performed on each trial from 5-35 Hz using Morlet wavelets (Morlet factor 7). Baseline correction was employed by subtracting the mean pre-movement (from −0.5 s to −0.02 s relative to the movement onset) from each trial. For two time windows of interest (peri-movement: from movement onset to movement offset; post-movement: from movement offset to 1 s after movement offset) β-power (13-30 Hz frequency range) was extracted for each trial, before averaging trial-wise β-power within the six bins. Repeated measures ANOVAs with bin as within-subjects factor were employed to analyse changes over time. To investigate associations between practice-induced change in behavioural measures and practice-induced change in β-power we computed Shepherd’s *pi* after outlier removal based on bootstrapping Mahalanobis distance from the bivariate mean^5^. Practice-induced change is defined as difference between the last and the first bin.

Hidden Markov Models (HMM) enable to describe the rich dynamics of electrophysiological data by inferring discrete functional states in a direct and data driven manner with millisecond precision, and without pre-specification of sliding window length (for a general introduction see^6^; for applications on MEG data see^7^^–^^9^; and for a tutorial see^10^). There are HMM variants for an overview), here we inferred the multivariate autoregressive ( MAR-) HMM^8^ (https://github.com/OHBA-analysis/HMM-MAR) on a single channel of data, corresponding to the 1^st^ PC of the group-concatenated single trials of the visuo-motor task data from twelve MEG planar gradiometers pairs covering the sensorimotor cortex. Since we are using the HMM-MAR on a single PC we refer to it as the *HMM-AR*. We used an HMM with K = 3 states (but see **Supplementary Fig.5** and **Supplementary Discussion**) and maximal order 5. To account for variations in the inference due to different initializations ten realizations were performed and the run with the lowest free energy chosen^7^. Inferred HMM states were characterized based on their time-course and spectral information, and out of these two measures reconstructed HMM-AR regularised time-frequency response^8^ (**Supplementary Fig.6**). Moreover, the relationship between single-trial time-course of grip force data and HMM state time-courses was quantified by means of Pearson’s correlation (**Fig.2**). Practice-induced changes in temporal characteristics of the inferred HMM states can be described using a range of metrics. Here, we focus on the fractional occupancy, lifetime (or dwell time) and transition probability (**Supplementary Fig. 4f-h**). Mirroring the procedure employed in the course of the Fourier-based, wavelet analysis, for the data acquired during the visuo-motor learning task HMM metrics were first averaged within each of the two time windows of interest (peri-movement: from movement onset to movement offset; post-movement: from movement offset to 1 s after movement offset) before averaging across trials within the six bins, to assess practice-related changes. Using the state time courses, local fractional occupancy (local FO) is defined as the average of the within-trial fraction of time spent in each state relative to the duration of the time window of interest, and fractional occupancy (FO) is the average across trials:

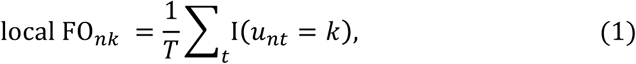

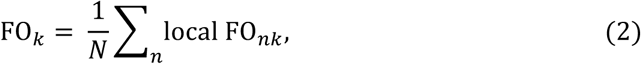

where *k* indexes the states, *u_t_* contains the value of the state time course at time point *t* for a single trial *n* within a time window of interest (*t*_0_ < *t* < *t*_1_), *T* is the length of *u_t_* in samples, *N* is the number of trials per individual, and function I in equation (1) returns 1 if the condition in its argument is true. Using the Viterbi path^12^ (which contains hard assignments instead of probabilities), local lifetimes (local LT) are defined as the average of the within-trial amount of time spent in each state before transitioning out of that state in a given time window, and lifetimes (LT) is the average across trials:

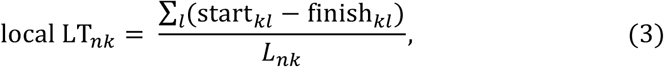

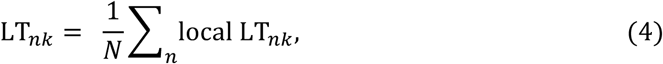

where *start_kl_*, and *finish_kl_*, respectively, denote the time points where the *l*-th visit to state *k* start and finish, and *L_nk_* is the number of occurrences for state *k* at trial *n*. The transition probabilities (TP) were computed from the state time courses, as described for example in^6,8,12^. Local TP describe the within-trial average of how likely it is to transition from one state to the same or another state in a given time window, and TP is the average across trials.

The **Supplementary Fig.8** depicts group averaged transition probability matrix. To investigate changes in fractional occupancy, lifetime and transition probabilities we performed repeated measures ANOVAs with bin as within-subjects factor. To investigate associations between practice-induced change in behavioural measures and practice-induced change in HMM metrics we computed Shepherd’s *pi* after outlier removal based on bootstrapping Mahalanobis distance from the bivariate mean^5^. Practice-induced change is defined as difference between the last and the first bin.

Focal learning-dependent transfer effects were evaluated by analysing pre-post changes in motor activation and rest. To ensure that we investigate the same property of the data, rather than global changes, we estimated the fractional occupancy, lifetime and transition probabilities from the motor activation and resting state data using the same model as for the visuo-motor learning data. In contrast to the visuo-motor learning data, motor activation and resting state data are not epoched, but continuous. Therefore, fractional occupancy is defined as fraction of time spent in each state relative to the total duration of the individual time course; lifetime is defined as the average amount of time spent in each state before transitioning out of that state over the whole time course; and transition probabilities describe the likelihood of transitioning from a given state to the same or another state over the whole time course. Fractional occupancy and lifetimes were calculated using equation (1), (3), respectively, whereby here *u_t_* denotes the whole state time course. Similarly, the transition probabilities where computed for the whole time course.

## Supplementary Results

Non-significant changes in the baseline suggest temporal specificity of the effects We have demonstrated significant learning-related changes in the β-HMM state, whereby the direction of change depend on the time window. This dissociation of effects suggests a movement-related temporal specificity in the practice-induced changes. To further explore the specificity of this potentially complex relationship, we additionally analysed practice-related changes in the β-HMM state during the time window that served as baseline for the Fourier-based analyses (-0.5 to -0.02 relative to movement onset). In contrast to the peri- and post-movement time windows, for the baseline time window motor practice did not result in any detectable changes (**Supplementary Table1**). Thus, our results suggest that motor practice leads to changes in specific, i.e. movement-relevant, time windows.

**Supplementary Table1:**
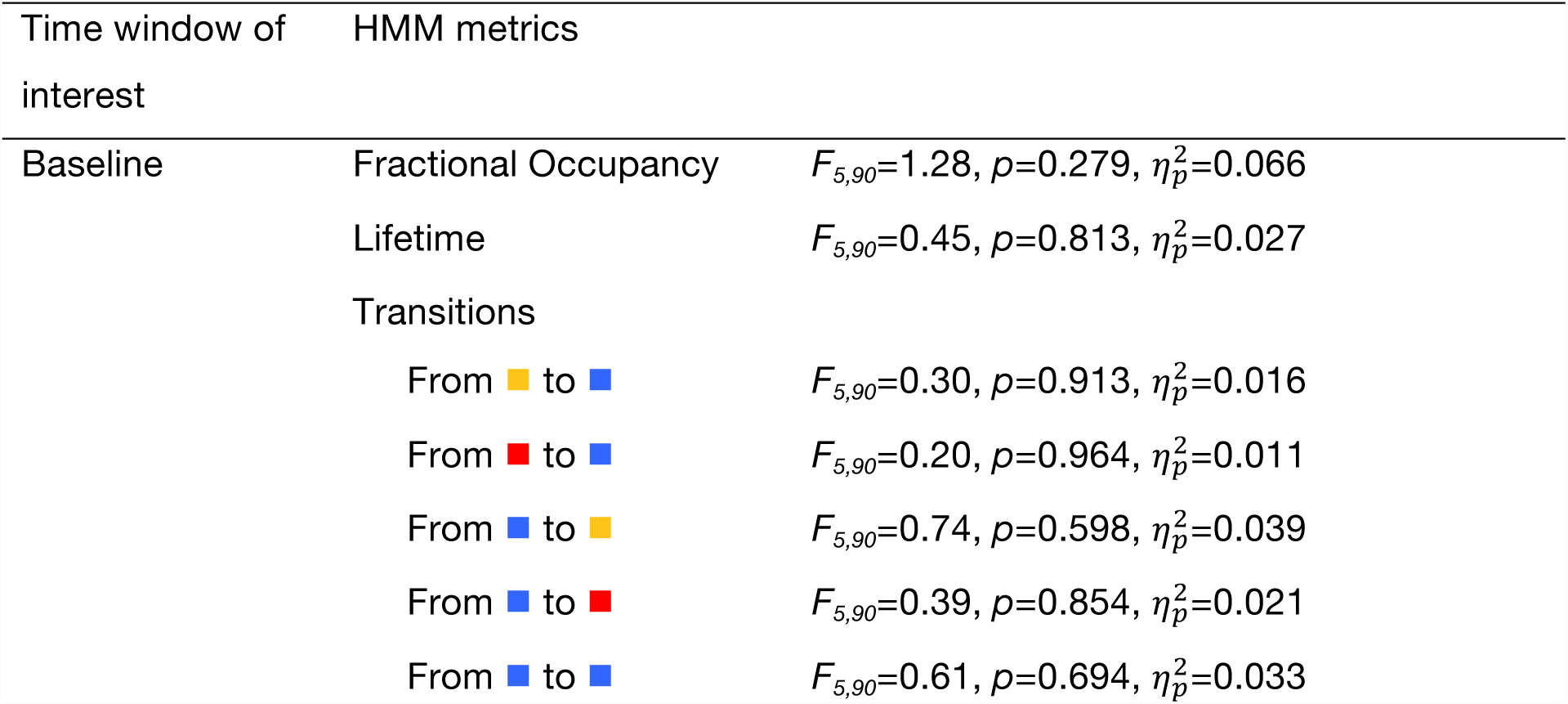
β-HMM state metrics for the baseline.

| Time window of interest | HMM metrics |  |
| --- | --- | --- |
| Baseline | Fractional Occupancy | $F_{5,90}=1.28, p=0.279, \eta_p^2=0.066$ |
| | Lifetime | $F_{5,90}=0.45, p=0.813, \eta_p^2=0.027$ |
|  | Transitions |  |
| | From to | $F_{5,90}=0.30, p=0.913, \eta_p^2=0.016$ |
| | From to | $F_{5,90}=0.20, p=0.964, \eta_p^2=0.011$ |
| | From to | $F_{5,90}=0.74, p=0.598, \eta_p^2=0.039$ |
| | From to | $F_{5,90}=0.39, p=0.854, \eta_p^2=0.021$ |
| | From to | $F_{5,90}=0.61, p=0.694, \eta_p^2=0.033$ |

### Absence of behaviourally relevant changes in the other two HMM states indicate spectral specificity of the effects

No systematic changes over the course of practice could be observed for the other two HMM states (yellow and red HMM state). Specifically, for the yellow state, neither peri-movement fractional occupancy, nor peri-movement lifetime were significantly modulated through practice (*F_5,90_*=0.36, *p*=0.874, 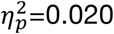; *F_5,90_*=1.11, *p*=0.361, 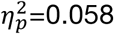; respectively; **Supplementary Fig.10**). For the same state, post-movement fractional occupancy decreased significantly over time (*F_5,90_*=3.46, *p*=0.007, 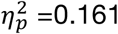), however, this change seems behaviourally not relevant (movement time: *pi_17_*=-0.09, *p*=1; accuracy: *pi_17_*=0.22, *p*=0.798), and lifetime remained unaltered (*F_5,90_*=1.00, *p*=0.422, 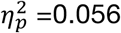). For the red state, perimovement fractional occupancy increased significantly over time (*F_5,90_*=4.06, *p*=0.002, 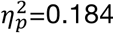), whereby this change did not correlate with behavioural changes (movement time: *pi_17_*=-0.37, *p*=0.310; accuracy: *pi_17_*=-0.30, *p*=0.478), and lifetime did not change significantly (*F_5,85_*=1.19, *p*=0.321, 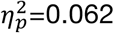). For the post-movement time window no significant alterations were found (fractional occupancy: *F_5,90_*=0.75, *p*=0.587, 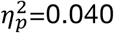; lifetime *F_5,85_*=0.68, *p*=0.637,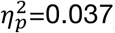). Systematic changes in transition probabilities were not observed for either of the two states (**Supplementary Fig.9**). In light of systematic learning-related changes in the β-HMM state and their absence in the other two states, it would appear that learning-related changes in the temporal dynamics of cortical sensorimotor β-activity are spectral specific.

### Absence of behaviourally relevant changes in a control ROI indicate spatial specificity of the effects

In order to test the spatial specificity of the effects we performed the same analyses on data from an anatomically defined *a priori* visual ROI covering the occipital pole. The three inferred HMM states show α-power spectral peaks, in the absence of other distinct spectral peaks (**Supplementary Fig.11a**). No clear movement-related changes in fractional occupancy could be observed (**Supplementary Fig.11 b**). Over the course of practice peri-movement fractional occupancy decreased for the pink state (*F_5,90_*=6.68, *p*<0.001, 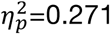) and increased for the black state ( *F_5,90_*=3.66, *p*=0.005, 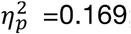; **Supplementary Fig.11c**). The absence of significant correlations between these changes in fractional occupancy and changes in performance (all *p-values*>0.1) suggest, however, that these changes are not behaviourally relevant for this task. The strong relationship between the opposing changes of peri-movement fractional occupancy between the black state and pink state (*pi_17_*=−0.88, *p*<0.001) rather indicate that over the course of training individuals swap their ‘preferred’ α-HMM state. No other changes in fractional occupancy, lifetime or transition probabilities were found for the peri- or post-movement time window (all *p*-values>0.1).

To investigate the aspect of spatial specificity even further we additionally interfered HMM metrics for the same visual ROI using the states derived from the sensorimotor ROI (**Supplementary Fig.12**). Similar to the analysis above, no clear movement-related changes in fractional occupancy could be observed. Over the course of practice peri-movement fractional occupancy decreased for the blue state (*F_5,90_*=6.31, *p*<0.001, 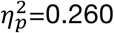) and increased for the red state (*F_5,90_*=4.33, *p*=0.001, 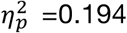; **Supplementary Fig.12c**). In addition, post-movement fractional occupancy increased for the red state _(_*F_5,90_*=2.54, *p*=0.034, 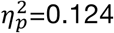). The absence of significant correlations between these changes in fractional occupancy and changes in performance (all *p-values*>0.1) suggest, however, that these changes are not behaviourally relevant for this task. No other changes in fractional occupancy, lifetime or transition probabilities were found for the peri- or post-movement time window (all *p*-values>0.1). Overall, HMM metrics estimated from a visual ROI using once three HMM states derived from the same visual ROI, and once the three HMM states derived from a sensorimotor ROI are very similar to one another, but seem not relevant for this task.

### Changes in post-movement sensorimotor β-activity and local accuracy

We found that practice-related changes in peri-movement, but not in post-movement, cortical sensorimotor β-activity were related to behavioural improvements. To further explore the relationship between post-movement cortical sensorimotor β-activity and accuracy we repeated our correlation analysis but focussed only on the last movement step of the sequential movement (i.e., gate 5). However, also in this case, no significant relationship between practice-related changes in accuracy and practice-related changes in post-movement β-activity could be observed (*pi_16_*=-0.18, *p*=1).

## Supplementary Discussion

### Number of States

One of the factors that need to be considered when employing HMM interference is the specification of the number of states. In the absence of a data driven approach to define the optimal number of states the final free energy of the HMM interference has often been considered. The final free energy of the HMM interference represents the lower bound of the so-called model evidence and represents a trade-off between model complexity and data likelihood (i.e. how well the model represents the data). In theory, it should be possible to select the optimal number of states by choosing the model with the lowest free energy. In practice however, it has been observed that the free energy increases almost monotonically^7^, which limits the practicability of this approach. Here the number of states was selected by choosing the model with the lowest free energy and the highest behavioural relevance. Behavioural relevance was defined as strongest relationship between time-course of grip force data and time-course of the three HMM states **Fig.2c** and **Supplementary Fig.5a**. Moreover, we repeated the original analysis for different numbers of states (K = 2, 4, 5, 6). The results, summarized in **Supplementary Fig.5**, demonstrate that the results were robust across different models. Interestingly, this is still true in the case of two β-HMM states (K = 5, 6), which indicates that the two β-HMM states represent similar properties in the data.

### Behavioural relevance of practice-related changes in post-movement sensorimotor β-activity

We could demonstrate that motor practice leads to faster and more accurate movements as well as more pronounced movement-related peri- and post-movement temporal dynamics of cortical sensorimotor β-activity. However, only practice-related changes in peri-movement, but not in post-movement, temporal dynamics of cortical sensorimotor β-activity were linked to behavioural improvements. This is contrary to findings from Tan and co-workers^13^, which is one of the very few studies investigating electrophysiological correlates of motor adaptation. Tan et al., report a trial-to-trial correlation between task accuracy and post-movement sensorimotor β-power obtained by conventional Fourier-based, wavelet analysis. However, there may be a number of reasons why it is not possible to directly compare their and our results. Tan et al., studied motor adaptation using a ballistic movement, whereas here we studied skill acquisition during a naturalistic sequential movement. Motor skill acquisition and motor adaptation are likely underpinned by distinct neural mechanisms^14^. In addition, while a ballistic movement and sequential movements with long inter-movement intervals comprise clear perimovement power decrease and post-movement power increase complexes, in sequential movements with short inter-movement intervals the post-movement power increase is interrupted or cancelled out by the pre- and peri-movement power decrease of the subsequent movement, leading to a phase of persistent power decrease^15^ (**Supplementary Fig.13**). Therefore, the post-movement power increase of a sequential movement might reflect mainly the last movement step of a sequential movement and not the total sequential movement. A repetition of the analysis, considering the accuracy for gate 5 only, did, however, not result in a significant correlation between practice-related change in accuracy and practice-related change in post-movement β-power (**Supplementary Results**). Without doubt, further electrophysiology studies are necessary to pinpoint the neural correlates of motor learning and motor adaptation in single and sequential movements.

## Footnotes

1 We use the term, peri-movement’ to refer to the time from movement onset to movement offset.

